# RB-TnSeq elucidates dicarboxylic acid specific catabolism in β-proteobacteria for improved plastic monomer upcycling

**DOI:** 10.1101/2025.02.21.639450

**Authors:** Allison N. Pearson, Julie M. Lynch, Cindy N. Ho, Graham A. Hudson, Jacob B. Roberts, Javier Menasalvas, Aaron A. Vilchez, Matthew R. Incha, Matthias Schmidt, Aindrila Mukhopadhyay, Adam M. Deutschbauer, Mitchell G. Thompson, Patrick M. Shih, Jay D. Keasling

**Author notes:** Corresponding authors : Jay D. Keasling,; Patrick M. Shih.

## Abstract

Dicarboxylic acids are key components of many polymers and plastics, making them a target for both engineered microbial degradation and sustainable bioproduction. In this study, we generated a comprehensive dataset of functional evidence for the genetic basis of dicarboxylic and fatty acid metabolism using randomly-barcoded transposon sequencing (RB-TnSeq). We identified four β-proteobacteria that displayed robust growth with dicarboxylic acid sole carbon sources and cultured their mutant libraries with dicarboxylic and fatty acids with carbon chain lengths from C3 to C12. The resulting fitness data suggested that dicarboxylic and fatty acid metabolisms are largely distinct, and different sets of β-oxidation genes are required for catabolizing dicarboxylic versus fatty acids of the same carbon chain lengths. Additionally, we identified transcriptional regulators and transporters with strong fitness phenotypes related to dicarboxylic acid utilization. In *Ralstonia sp.* UNC404CL21Col (*R. CL21*), we deleted two transcriptional repressors to improve its utilization of short chain dicarboxylic acids. We exploited the diacid-utilizing catabolism of *R. CL21* to upcycle a mock mixture of the dicarboxylic acids produced when polyethylene is oxidized. After introducing a heterologous indigoidine production pathway, this engineered *Rastonia* produced 0.56 ± 0.02 g/L indigoidine from a mixture of dicarboxylic acids as a carbon source, demonstrating the potential of *R. CL21* to upcycle plastics waste to products derived from tricarboxylic acid (TCA) cycle intermediates.

**Importance:** Upcycling the carbon in plastic wastes to value-added products is a promising approach to address the plastics waste and climate crises, and dicarboxylic acid metabolism is an important facet of several approaches. Improving our understanding of the genetic basis of this metabolism has the potential to uncover new enzymes and genetic parts for engineered pathways involving dicarboxylic acids. Our dataset is the most comprehensive interrogation of dicarboxylic acid catabolism to date, and this work will be of utility to researchers interested in both plastics bioproduction and upcycling applications.

## Introduction

Dicarboxylic acids are foundational industrial chemicals, serving as precursors for numerous products, including plastics and other polymeric materials (1, 2). Conversely, they are also one of the products of several plastics degradation methods (3–6). With roughly 8.3 billion metric tons of plastics produced between the industry’s post-WWII inception and 2017 – often from fossil fuel-derived precursors – there is a need to establish both alternative disposal methods for existing plastics and sustainable production of novel, biodegradable plastics (7). Therefore, microbial upcycling of plastics into value-added products and microbially-produced bioplastics are both critical areas of current metabolic engineering research (1, 8, 9).

To reduce our reliance on petroleum-based polymers, there have been many recent efforts in engineering the bioproduction of dicarboxylic acids (1, 2, 10–12). Approaches include ⍵-oxidation of accumulated fatty acids (13), diverting flux from native pathways (e.g., biotin biosynthesis) (14), engineered pathways utilizing polyketide synthases (15), and reversed β-oxidation (16, 17), amongst others. Many of these engineered pathways require heterologous expression of β-oxidation enzymes and transporters to function efficiently, highlighting the necessity of genetic knowledge pertaining to the natural metabolism of the target dicarboxylic acid of interest. This need to understand the genetic basis of dicarboxylate metabolism is also apparent in current plastics upcycling research. In recent years, advances in the breakdown of plastics have increased interest in microbial plastic upcycling (8, 9, 18). Plastics can be degraded into oligomers and monomers suitable for microbial feedstocks with methods such as chemical treatments (4, 5, 19), enzymatic treatments (20), metal catalysts (6), and whole-cell biocatalysis (21, 22). For some plastics, many of these degradation methods result in dicarboxylic acids of varying chain lengths (3–6). Microbial upcycling of dicarboxylic acids has been demonstrated – for example, *Pseudomonas putida* has been engineered to produce polyhydroxyalkanoates, cis-cis muconate, and β-ketoadipate from dicarboxylic acids (3, 23, 24). In the aforementioned examples, it was first necessary to engineer the efficient consumption of dicarboxylic acids in the host organism (3, 23, 24).

It is well-established that microbes metabolize dicarboxylic acids through β-oxidation, yet it is not always obvious which β-oxidation pathways can act on dicarboxylic acids. β-oxidation genes can be specific for certain substrates; for example, *P. putida* can natively consume medium-chain length fatty acids (FA) but cannot consume dicarboxylic acids (DA) of the same chain lengths. Heterologous expression of the adipate β-oxidation genes *dcaAKIJP* from *Acinetobacter baylyi* was required for dicarboxylic acid consumption (23). This highlights the importance of obtaining functional evidence in order to characterize dicarboxylic acid metabolisms. Although dicarboxylic acid β-oxidation pathways have been identified in several organisms (25–27), increasing the number of genes with functional evidence of dicarboxylic acids activity has the potential to reveal new genetic tools for improving both dicarboxylic acids bioproduction and upcycling.

Randomly barcoded transposon sequencing (RB-Tn-Seq) allows rapid genome-wide profiling of individual gene fitness under a selective condition in a high-throughput manner. In this method, a library of transposon mutants – each carrying a DNA barcode that is mapped to the gene it disrupts – is created. This library is exposed to a selective condition that will result in the enrichment or depletion of some barcoded mutants, depending on whether disruption of that gene was beneficial, neutral, or detrimental in the selective condition. RB-TnSeq has been used to study the metabolism of industrially relevant compounds in *P. putida* KT2440, including fatty acid, alcohol, and ⍵-aminocarboxylic acids, enabling the identification of genes required for their utilization as sole carbon and/or nitrogen sources. This work also revealed some of the genetic elements responsible for the regulation of these metabolisms, which have the potential to be leveraged as genetic tools in future work (28–30).

Here, we leveraged RB-TnSeq to interrogate the genetic basis of dicarboxylic acid catabolism and investigate the potential for hosts for plastics upcycling. We grew 12 organisms for which there exists an RB-TnSeq library on fatty acids and dicarboxylic acids of various carbon chain lengths and identified four strains of β-proteobacteria that demonstrate promising growth. RB-TnSeq fitness data revealed genes involved in dicarboxylic acid catabolism, along with transporters and regulators genes related to that metabolism, in all four β-proteobacteria. Based on these fitness data, a double knockout strain of *Ralstonia sp.* UNC404CL21Col (*R. CL21*) was developed to enable utilization of additional dicarboxylic acids. We also evaluated *R. CL21* as a potential host for plastics upcycling and demonstrated high titer production of indigoidine from a mixture of dicarboxylic acids as a proof of concept.

## Results and Discussion

### Screen for dicarboxylate metabolizing organisms

To identify dicarboxylic acid catabolic pathways, we began with bacteria for which RB-TnSeq libraries already existed. Twelve of these bacteria (*Burkholderia phytofirmans* PsJN, *Cupriavidus baseilensis* FW507-4G11, *Dyella japonica* UNC79MFTsu3.2, *Escherichia coli* BW25113, *Herbaspirillum seropedicae* SmR1, *Klebsiella michiganesis* M5aI, *Paraburkholderia bryophila* 376MFSha3.1, *Pedobacter sp.* GW460-11-11-14-LB5, *Pseudomonas fluorescens* FW300-N2C3, *Pseudomonas simiae* WCS417, *Ralstonia sp.* UNC404CL21Col, and *Sinorhizobium meliloti* 1021) could grow in MOPS-buffered minimal medium with succinate and spanned the *Bacteroidota* and *Pseudomonadota* phyla (Figure 1A). We tested the growth of these bacteria with fatty and dicarboxylic acids with carbon chain lengths of 4-10 (C4-C10) as sole carbon sources (Figure 1B). The pseudomonads grew particularly well on the medium chain fatty acids, which was consistent with previous research (30). However, only the β-proteobacteria *Burkholderia phytofirmans* PsJN*, Cupriavidus basilensis* FW507-4G11*, Paraburkholderia bryophila* 376MFSha3.1, and *Ralstonia sp.* UNC404CL21Col grew on the medium chain dicarboxylic acids.

**Figure 1:**
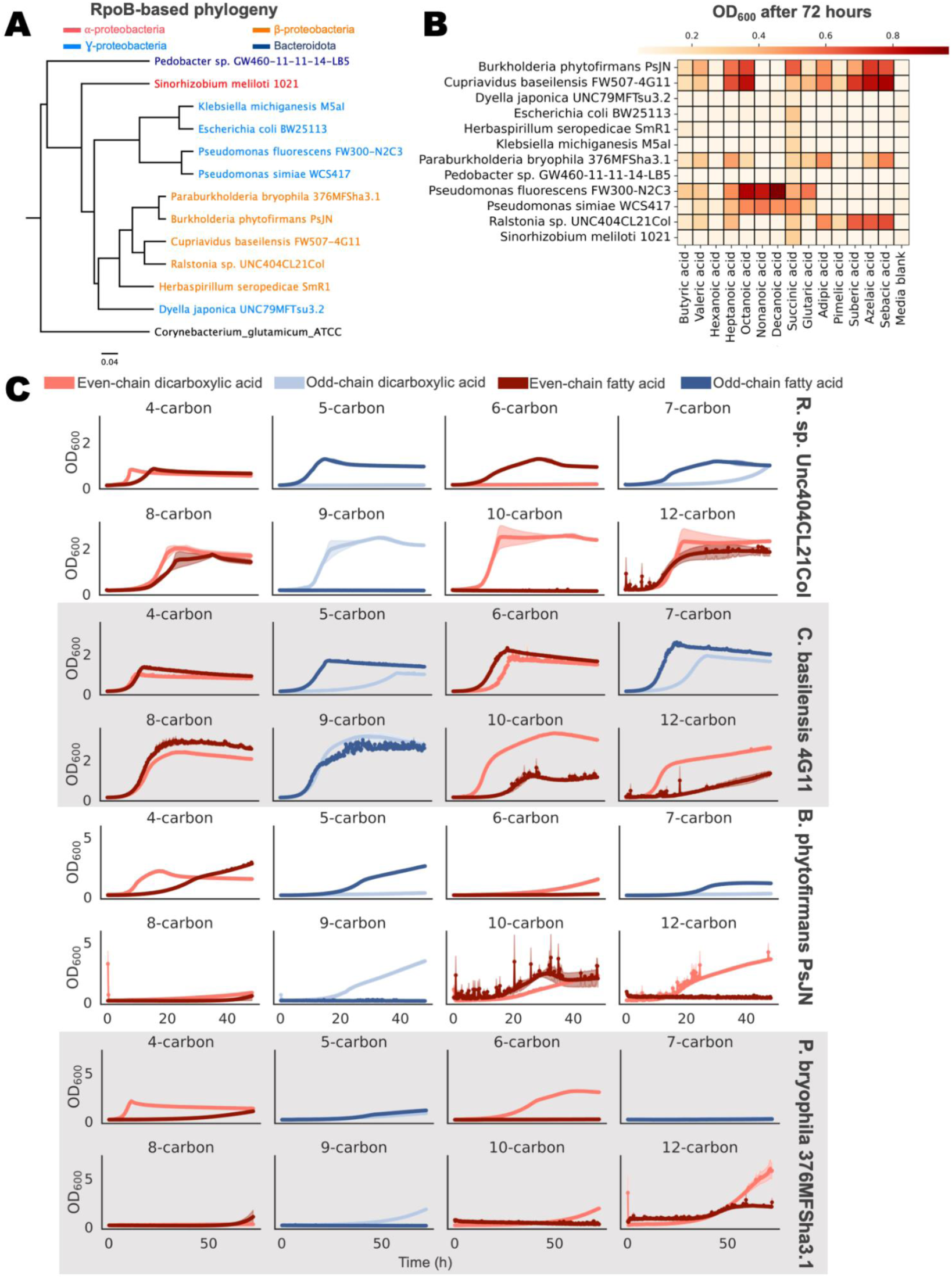
Identifying candidate organisms that can utilize dicarboxylic and fatty acids as sole carbon sources. A) Phylogenetic tree of candidate organisms based on RpoB sequence. Taxa are colored according to their phylum/subphylum (Bacteroidetes = dark blue, ɑ-proteobacteria = red, β-proteobacteria = orange, ɣ-proteobacteria = light blue). B) Optical density at 600 nm after 72 hours for growth of 12 organisms on 7 fatty acids and 7 dicarboxylic acids in MOPS minimal medium (n=3). C) Growth of *R. CL21*, *C. basilensis*, *B. phytofirmans,* and *P. bryophila* on dicarboxylic and fatty acid carbon sources, measured by OD_600_ (n=3, error = 95% confidence interval).

Kinetic growth assays of these four organisms revealed differences in growth between dicarboxylic versus fatty acid carbon sources (Figure 1C). For instance, *R. CL21* was unable to grow with glutarate (C5-DA) as a sole carbon source and demonstrated limited growth with the shorter chain length dicarboxylic acids (adipic (C6) and pimelic (C7) acids), even though the strain displayed significant growth with the equivalent chain length of fatty acids. This trend was reversed for the C9 and C10 chain length fatty and dicarboxylic acids, with *R. CL21* unable to utilize the fatty acids but demonstrating robust growth on the dicarboxylic acids. Growth was comparable between strains grown with C8 and C12 fatty and dicarboxylic acid carbon sources. Similar trends were present in the other three organisms, which tended to grow more robustly with C8+ dicarboxylic acids than with C8+ fatty acids or shorter chain length dicarboxylic acids. The strong differences between dicarboxylic and fatty acid growth in these β-proteobacteria therefore presented a rich scenario for functional analysis with RB-TnSeq.

### Investigating dicarboxylic acid metabolism with RB-TnSeq

RB-TnSeq experiments were performed in duplicate for the four β-proteobacteria with acetate and fatty/dicarboxylic acids with chain lengths from C3-10 and C12 as the sole source of carbon in MOPS-buffered minimal medium. As a control to determine which phenotypes were specific for fatty/dicarboxylic acids, we also conducted D-glucose carbon source experiments for three of the four libraries. Since *C. basilensis* cannot effectively utilize D-glucose as a sole carbon source, D-glucose was replaced with DL-lactate as the control for that library. Finally, we also tested protocatechuic acid, since the ortho-cleavage pathway of its metabolism results in the dicarboxylic acid β-keto-adipate (Supplementary Table 1) (27, 31). The complete dataset is publicly available on the Fitness Browser (fit.genomics.lbl.gov) (32). Across all four organisms, we identified 1,349 genes for which there was a significant and fatty/dicarboxylic acid specific phenotype with at least one of our tested carbon source conditions. We define a significant phenotype as a gene having an |fitness score| > 1 and an |t-score| > 4, while we define a fatty/dicarboxylic acid specific phenotype as a gene having a significant fitness score in at least one condition of interest but no significant score when D-glucose (lactate for *C. basilensis*) was the provided carbon source.

To visualize the fitness data across all four organisms, we used the manifold learning method t-stochastic neighbor embedding (t-sne) for dimensionality reduction based on the fitness profile of each of the 1,349 genes with significant and specific phenotypes (Figure 2A). Although there were not many fully distinct clusters, there were clear regions where DA and FA genes are largely segregated (Figure 2A, left). The regions of high overlap and limited number of distinct clusters were not surprising given the iterative nature of β-oxidation; many genes had significant fitness phenotypes over a range of conditions, so very few distinct clusters emerged. However, in some cases the clustering did follow patterns with clear biological significance. For example, the group centered around coordinates (75, 20), consisted largely of predicted propionate utilization and methylcitrate cycle genes, corroborated by their significant phenotypes with odd chain fatty acid carbon source conditions. This analysis can be browsed interactively, with a display of gene name, annotation, and hyperlinks to the fitness browser, in Supplemental File 1.

**Figure 2:**
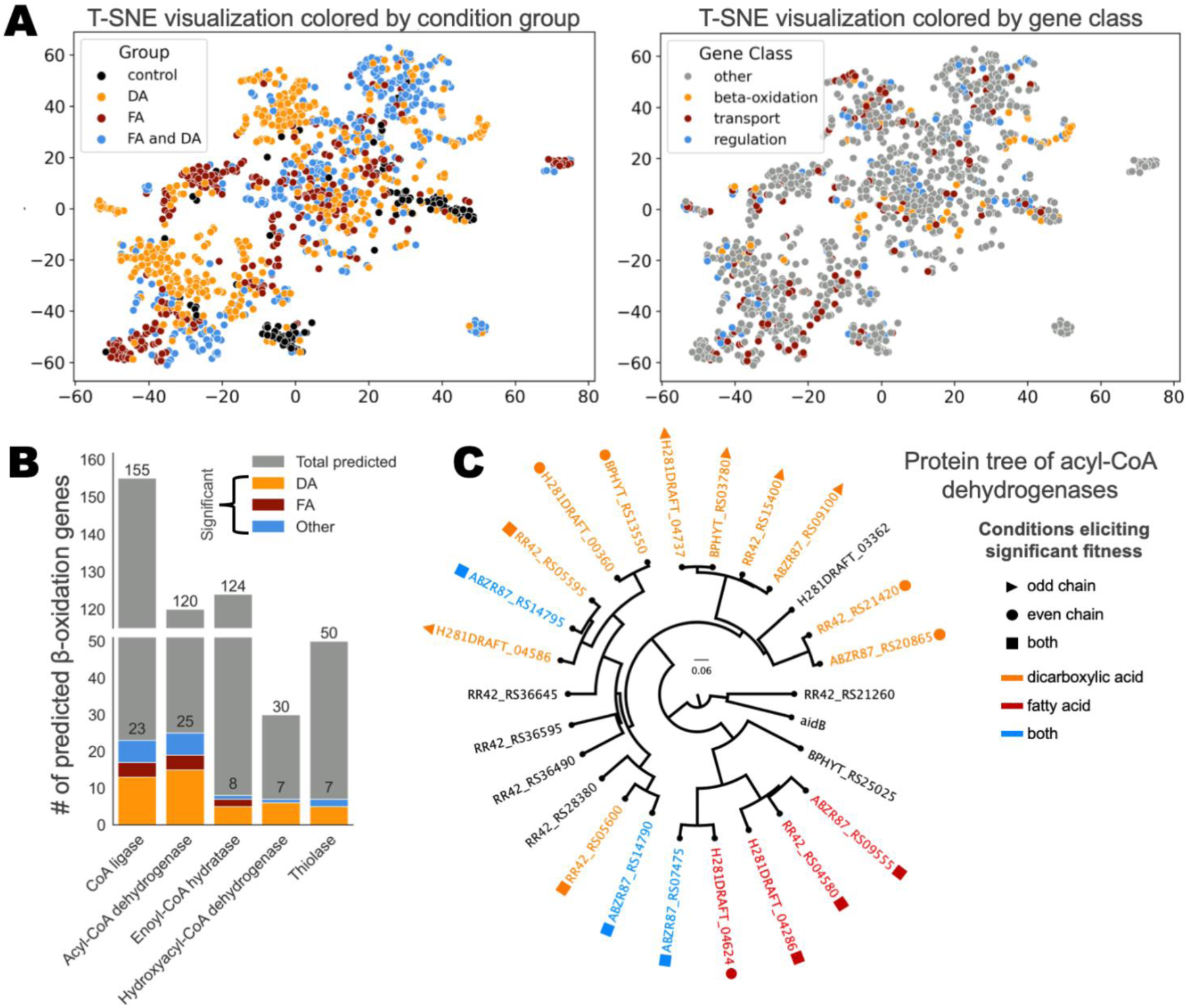
RB-TnSeq reveals genes involved in dicarboxylic acid and fatty acid metabolism. A) Genes with a significant fitness phenotype across four β-proteobacteria, plotted by dimensionality reduction of their fitness profile via t-sne. Subplots vary only by the metric used to determine the color of the plotted genes, either if they have phenotypes for dicarboxylic or fatty acids, shorter or longer carbon chain lengths, or even or odd carbon chain lengths. B) Total number of predicted β-oxidation genes across all four β-proteobacteria. β-oxidation genes with significant fitness are colored according to the conditions that elicited significant phenotypes: dicarboxylic acids only (orange), fatty acids only (red), or fatty/dicarboxylic acids and/or the control conditions (blue). Genes encoding β-oxidation enzymes were predicted by Pfam, using HMM (Supplementary Tables 6-10). C) Phylogenetic tree of acyl-CoA dehydrogenase genes with significant fitness phenotypes. Genes with a negative fitness phenotype are colored according to which conditions elicit this phenotype (orange = dicarboxylic acid, red = fatty acid, and blue = both) and have a shape indication whether they have significant phenotypes for odd (triangle), even (circle), or both types of (square) carbon chains. The protein tree was constructed using MUSCLE with the super5 algorithm, and the UPGMA algorithm for tree construction. The protein sequence for *E. coli* AidB was included as an outgroup to confirm the rooting of the tree.

More importantly, we found that most genes had significant phenotypes on either DAs (37.1%) or FAs (23.3%), and only 30.5% of genes were significant for both (Figure 2A, left). Additionally, only 24.7% of significant genes were predicted to be directly involved in transport, regulation, and β-oxidation; the other 1,016 significant genes were not predicted to fall into one of these categories (Figure 2A, right). Altogether, this suggests that DA and FA metabolisms are largely genetically distinct and elicit a physiological response beyond their direct β-oxidation.

We predicted which genes across the four organisms encode β-oxidation enzymes by Pfam using HMMER (Supplementary Tables 6-10) and subsequently identified which β-oxidation genes had significant and specific fitness in one or more fatty/dicarboxylic acid conditions. Across all four genomes there were 155 predicted acyl-CoA ligases (23 significant/specific), 120 predicted acyl-CoA dehydrogenases (25 significant/specific), 124 enoyl-CoA hydratases (8 significant/specific), 30 3-hydroxyacyl-CoA dehydrogenases (7 significant/specific), and 50 thiolases (7 significant/specific) (Figure 2B, Supplementary Tables 6-10). Due to the presence of bifunctional enoyl-CoA hydratases/3-hydroxyacyl-CoA dehydrogenases, there were some duplicate genes between these two categories. As with the dataset as a whole, there were more β-oxidation genes that had specific phenotypes for either dicarboxylic or fatty acids that there were for both, reaffirming the notion that these metabolisms are separate and agreeing with previous work in the β-proteobacterium *Cupriavidus necator* H16 (26). The greatest number of significant predicted β-oxidation genes (20+) fell into the CoA-ligase and acyl-CoA dehydrogenase categories. This suggests that these steps of β-oxidation may be the most selective when it comes to substrate specificity. The later β-oxidation steps had less than half of the amount of significant genes per step seen in the first two steps. This implies that these organisms have more functional redundancy and less specificity in their enoyl-CoA hydratases/3-hydroxyacyl-CoA dehydrogenases and thiolases. This is especially true for the fatty acids, for which there are only two specific β-oxidation genes for steps after the oxidation of the fatty acyl-CoA – the homologous enoyl-CoA hydratases H281DRAFT_01199 and ABZR87_17485 (81% similarity).

To explore how the evolution of these genes might be related to their function, we constructed a phylogenetic tree of the significant/specific acyl-CoA dehydrogenases protein sequences, for which there were the greatest number of significant/specific genes (Figure 2C, Figure S2A). As an outgroup, we selected an *E. coli* gene for AidB, a conserved SOS-response protein that shares structural similarity and a common ancestor with acyl-CoA dehydrogenases (33, 34). One of our predicted acyl-CoA dehydrogenases in *C. basilensis*, RR42_RS21260, formed a clade with the AidB sequence, indicating that it is likely an AidB homolog and not a conventional acyl-CoA dehydrogenase. RR42_RS21260 only had a significant phenotype in the adipic acid carbon source condition, in which its disruption was beneficial with a positive score of 1.5 (Figure S2A). To visualize the relationship between phylogenetic and functional relatedness, proteins in the tree were classified according to whether their genes had significant negative phenotypes in fatty acid or dicarboxylic acid conditions and conditions with even or odd numbers of carbon (Figure 1C). The AidB homolog and five true acyl-CoA dehydrogenases that only had positive significant phenotypes were not considered. Two clades emerged in the phylogenetic tree, one that included mostly acyl-CoA dehydrogenases that appear to act on fatty acid substrates, and another that includes those that mostly seem to act on dicarboxylic acid substrates. In fact, the two genes with a significant negative fitness phenotype for both types of substrates in the ‘dicarboxylic acid clade’, ABZR87_RS14790 and ABZR87_RS14795, only had a significant fitness defect with one fatty acid substrate, heptanoic acid. At an average of -1.1, their fitness defect with heptanoic acid was much weaker than their defect for dicarboxylic acid substrates suberic acid (-5.0), azelaic acid (-4.7), sebacic acid (-6.0), and dodecanedioic acid (-5.2) (Figure 1C, Figure S2). Interestingly, these two predicted acyl-CoA dehydrogenase genes appear in the same operon and have very similar fitness profiles, and homologs with similar fitness profiles and synteny are present in the other three organisms. ABZR87_RS14790 and ABZR87_RS14795 have high protein sequence similarity to the pimeloyl-CoA dehydrogenase PimCD from *Rhodopseudomonas palustris* (74% similarity, 100% coverage) and the sterol acyl-CoA dehydrogenase ChsE12 from *Mycobacterium tuberculosis* (43% similarity, 99% coverage) (27, 35–37). The latter of these has been demonstrated to function as an α_2_β_2_ heterotetramer rather than the homotetramer or homodimer characteristic of acyl-CoA dehydrogenases (35) (37). It is possible that this dicarboxylic acid acyl-CoA dehydrogenase may also function as a heterotetramer.

Within the dicarboxylic acid clade of acyl-CoA dehydrogenases, we also see a group – with a representative from each organism – that has fitness phenotypes on odd-chain dicarboxylic acids (ABZ87_RS09100, RR42_RS15400, H281DRAFT_04737, and BPHYT_RS_03780) (Figure 2C, Figure S2A). All of these genes share more than 84% identity over 100% coverage with the glutaryl-CoA dehydrogenase PP_0158 from the decarboxylative glutarate degradation pathway in *P. putida* KT2440, and likely catalyze the oxidative decarboxylation of glutaryl-CoA into crotonyl-CoA and CO_2_ (38). This is supported by their phenotypes with the pimelic (C7DA) and azelaic (C9DA) acid conditions, which after successive rounds of β-oxidation result in glutaryl-CoA (Figure S2A).

The one outlier in the fatty acid specific clade of acyl-coA dehydrogenases, ABZR87_RS07475, appears to be promiscuous, with significant fitness defects with protocatechuate (-2.1), fatty acid (-1.7 for heptanoic acid, -2.5 for octanoic acid), and dicarboxylic acid (-3.1 for sebacic acid, -1.1 for azelaic acid) carbon sources. However, the remaining four members of the clade had significant fitness defects only in fatty acid carbon source conditions. Overall, this phylogenetic tree reinforces the idea that the ability to metabolize dicarboxylic acids is often distinct from the ability to metabolize fatty acids, and requires the evolution of separate β-oxidation genes.

Finally, this analysis demonstrates which acyl-CoA dehydrogenase genes are homologous across the four organisms and could be useful to researchers endeavoring to utilize these genes to engineer dicarboxylic acid metabolism in other organisms. Towards that end, Figure S2B-C shows a phylogenetic tree of the CoA ligases and a heatmap of their fitness, and Supplemental Table 2 displays all β-oxidation genes with highly similar sequences to each other.

### Dicarboxylic acid transport and regulatory genes revealed by RB-TnSeq

In addition to the genes directly involved in β-oxidation, we also identified transcriptional regulators and transporters required for dicarboxylic acid catabolism. Transcriptional regulators are of interest because they can be utilized for dynamic gene regulation or as potential biosensors, while transport proteins can be utilized to control the import/export of dicarboxylic acids essential for engineering both dicarboxylic acid consumption and bioproduction. For example, heterologous expression of the *A. baylyi* dicarboxylate uptake genes *dcaKP* were necessary for engineered dicarboxylic acid consumption in *P. putida (23)*. Across the four β-proteobacteria, we found 3,439 predicted transport genes and 1,468 predicted regulatory genes, of which 173 and 95 had significant fitness phenotypes, respectively (Figure 3A). It should be noted that many transporters and regulators function as heteromeric complexes, and the number of genes does not directly correlate with the number of distinct systems. *R. CL21* had the greatest number of significant predicted transport and regulator genes, at 72 and 31, respectively, over double the amount predicted in *B. phytofirmans* (Figure 3B). Note that these numbers may be affected by the number of conditions in which the different libraries passed metric; the *R. CL21* library passed metric in 17 distinct experimental conditions while the *B. phytofirmans* library only resulted in high quality data in 10 conditions (Supplementary Table 1). We also classified transporter and regulator genes by whether they had significant phenotypes with dicarboxylic acid, fatty acid, or both condition classes. Consistent with our global analysis, we found that a large portion of significant transport and regulatory genes had significant fitness with either dicarboxylic acid or fatty acid conditions, not both (Figure 3C). The fitness data for all predicted transport and regulatory genes with significant phenotypes can be found in Figures S3-S6.

**Figure 3:**
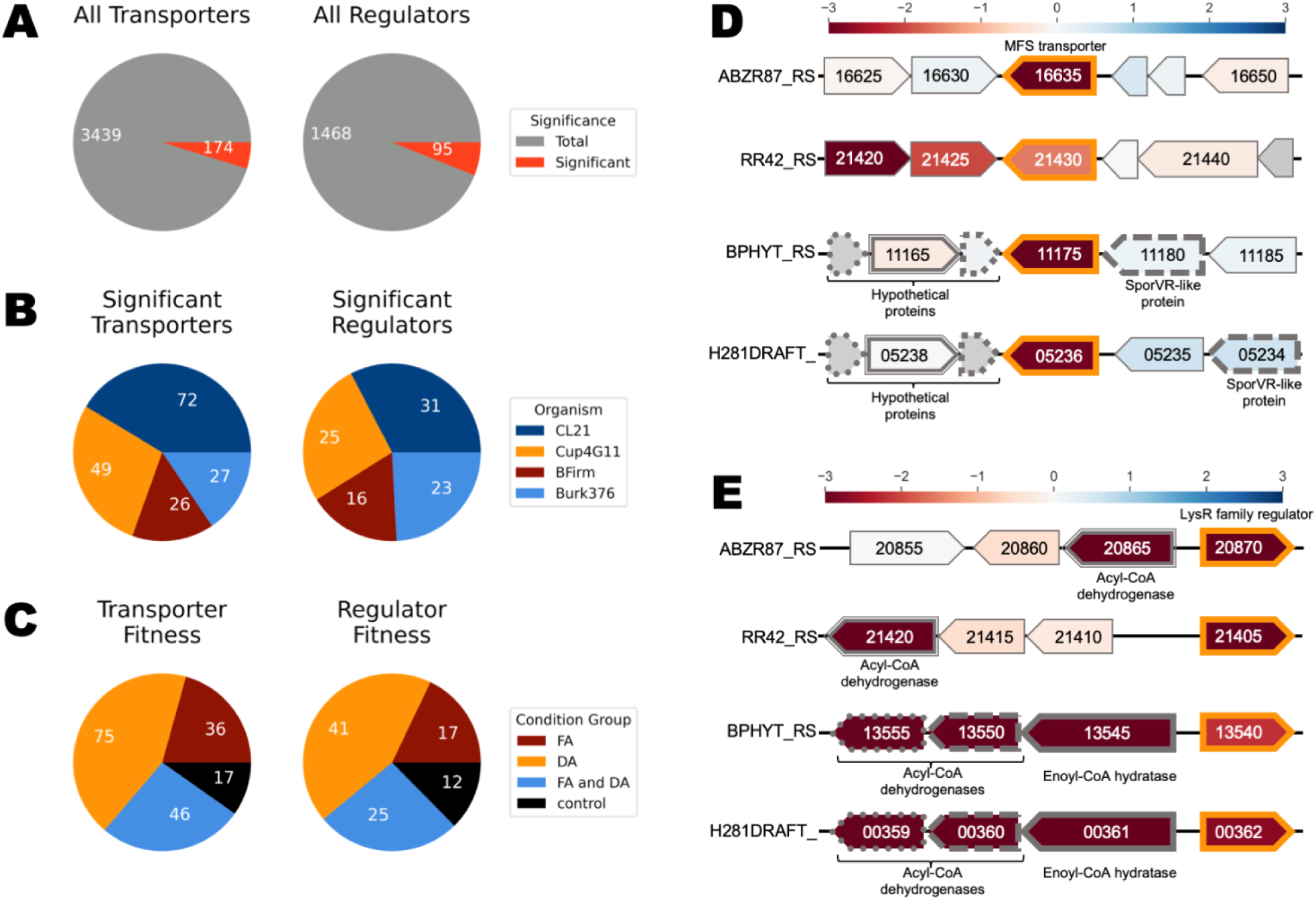
RB-TnSeq reveals transport and regulatory genes with significant and specific phenotypes. A) Total number of predicted transport and regulatory genes. Transport genes were identified using TransAAP, and regulatory genes were identified via the Pfams shown in Supplementary Table 3 with an e-value cutoff of 1e-20. Significant genes were defined as genes that had |fitness| > 1 and |t-score| > 4 with at least one of the experimental conditions, and did not elicit a significant phenotype in their glucose/lactate control. B) Breakdown of significant transporters and regulators by organism. C) Breakdown of significant transporters and regulators by condition eliciting significant phenotypes – either fatty acids (FA), dicarboxylic acids (DA), both (FA and DA), or controls. D) A synteny plot of a conserved adipic acid transporter in all four organisms (outlined in orange). Genes with > 70% similarity are indicated by matching outline patterns and their predicted function indicated. Gene sizes are approximate. Gene color corresponds to fitness in the adipic acid condition, with gray indicating genes for which there was no fitness data. E) A synteny plot of a conserved LysR family regulator in all four organisms (outlined in orange). Genes with > 70% similarity are indicated by matching outline patterns and their predicted function indicated. Gene sizes are approximate. Gene color corresponds to fitness in the adipic acid condition.

There were numerous significant transport systems that were conserved across the four organisms. For example, each had a homolog of a predicted α-ketoglutarate permease with a significant negative fitness phenotype for adipic acid (ABZR87_RS16635, RR42_RS21430, BPHYT_RS11175, H281DRAFT_05236, >82% similarity) (Figure 3D). The fitness data indicated that, with the exception of the *P. bryophila* homolog, these transporters may function specifically for adipic acid transport (Figures S3A-S6A). The *P. bryophila* homolog, H281DRAFT_05236, also had significant phenotypes in the pimelic and suberic acid conditions, suggesting that it could act more broadly on medium chain length dicarboxylic acids (Figure S6B). Despite the apparent conservation of function and high level of protein similarity across the four organisms, it appeared that the genomic context of this transporter was only conserved between *P. bryophila* and *B. phytofirmans* (Figure 3D).

Another notable highlight of the transport data was the presence of two putative pimelic acid-specific transport systems in *R. CL21*. All the genes in the predicted ABC transporter ABZR87_RS12600-20 had strong negative fitness specific to pimelate (C7DA) (Figure S3A). *C. basilensis* appears to have a homologous transporter, RR42_RS11130-RR42_RS11150, with the least similar subunit still sharing a protein sequence similarity of 70% identity with 94% coverage with ABZR87_RS12600-20. The transporter in *C. basilensis* appears less specific than the one in *R. CL21*, with negative phenotypes in both pimelic and adipic acids (Figure S4A). ABZR87_RS04195-215, also a predicted ABC transporter, had significant negative fitness phenotypes on pimelic acid as well (Figure S3A). The homologous operons from the other three organisms (70+% identity, 90+% coverage) did not have a phenotype in pimelic acid, although it should be noted that the *B. phytofirmans* library did not pass metric in the pimelic acid condition.

In *R. CL21* and *C. basilensis*, these pimelic acid transporters did not appear to act on the C7 fatty acid, heptanoate. Instead, a different predicted ABC transporter had significant fitness defects in the C7 and C8 fatty acid conditions (Figure S3A, S4A). The genes for this transporter, ABZR87_RS02545-50 and RR42_RS18960-70, are organized in an operon and share at least 75% identity over 84% coverage. Interestingly, although the strongest fitness phenotypes for these genes are in the fatty acid conditions, there is also a weak phenotype for this transporter in the dodecanedioic acid (C12DA) condition in both organisms, suggesting that it may be promiscuous.

Amongst the predicted regulators with significant fitness, there were also several notable genes (Figures S3B-S6B). In *R. CL21*, the LysR family regulator ABZR87_RS20870 had a significant and specific negative fitness phenotype on all of the C6-C12 dicarboxylic acids, while its nearest homologs in *C. basilensis* and *B. phytofirmans,* RR42_RS21405 and BPHYT_RS13540, only had a phenotype for some of the even chain length dicarboxylic acids (Figures 3E, S3B, S4B, S5B). In *P. bryophila*, the nearest homolog to this LysR family regulator, H281DRAFT_05209, appeared to be specific for adipic acid (Figure 3E, S6B). The homologs in *B. phytofirmans/P. bryophila* are more similar to each other (95%) than they are to those in *R. CL21* and *C. basilensis* (67-72 % similarity), and share the same genomic context with adjacent predicted enoyl-CoA hydratase and acyl-CoA dehydrogenase genes (Figure 3E). These genes have similar fitness profiles and scores as their neighboring LysR family regulators, suggesting that they are likely part of its regulon (Figure 3E). However, this regulator does not have the same genomic context in *R. CL21/C. basilensis*. There was a predicted acyl-CoA dehydrogenase near the regulator in these organisms, but it was not homologous to the pairs found in *B. phytofirmans/P. bryophila* (Figure 3E). Instead, the *R. CL21*/*C. basilensis* homologs to the pairs of acyl-CoA dehydrogenases found in *B. phytofirmans/P. bryophila,* with >58% similarity, were located elsewhere in the genome and had fitness profiles that differed slightly from that of the LysR family in *R. CL21/C. basilensis* (Figures 3E, S2A,S3A, S4A). This suggests that the regulons of this LysR family regulator may not be completely conserved across the organisms. *C. basilensis* also had another predicted transcription factor, RR42_RS20375, with significant fitness specific to dicarboxylic acids, with negative phenotypes on glutaric, adipic, pimelic, and suberic acids (Figure S4B). There was no homolog of RR42_RS20375 with significant fitness in any of the other organisms.

To explore the relationship between sequence similarity and fitness similarity in the regulators and transporters, we conducted a genotypic and phenotypic comparison of every gene-gene pair. Using pairwise BLASTp and a Pearson correlation of fitness profiles between all possible pairs of transport/regulator protein sequences with significant fitness phenotypes, gene pairs were plotted according to their fitness correlation and percent identity (Figures S7). As expected, among protein sequence pairs with high sequence correlation, there was a concentration of high positive fitness correlation. This region likely correlates to homologs with conserved functions, for example, the aforementioned conserved α-ketoglutarate permease. However, there were also gene pairs with fitness correlations near zero but high similarity, corresponding to homologs with divergent functions. Gene pairs with negative fitness correlation indicated reversed phenotypes and could suggest opposite functions or differences in physiological states, which is especially interesting in the case of high sequence similarity. An example of this is the predicted LysR family regulator, which had negative fitness correlations (∼-0.5) but high similarity (∼80%). This regulator is predicted to be a homolog of OxyR, which positively regulates microbial responses to oxidative stress in response to conditions like hydrogen peroxide (39). The homologous LysR family regulators BPHYT_RS03470 and ABZR87_RS01255 have a positive fitness phenotype with protocatechuate, while RR42_RS17385 has a negative phenotype with protocatechuate. This could indicate a difference in oxidative stress between the species in the presence of protocatechuate. Interactive versions of these visualizations can be found in Supplemental Files 2 and 3.

### β-oxidation in Ralstonia CL21

We next decided to take a closer look at the catabolism of dicarboxylic acids in *R. CL21*, which, out of the four libraries tested, resulted in fitness data with the greatest number of carbon sources and could have potential as a new chassis for dicarboxylic acid upcycling. We identified the predicted β-oxidation, glyoxylate shunt, and methylcitrate cycle genes that had a significant and specific phenotype in *R. CL21* (Figure 4). Just as we observed across all four organisms, many genes appeared dedicated to either FA or DA metabolism, and many genes have very strict chain length specificity, which agrees with previous work (26). The same analysis and data visualization was also performed on the other three β-proteobacteria, and can be found in the Supplemental materials (Figures S8-10).

**Figure 4:**
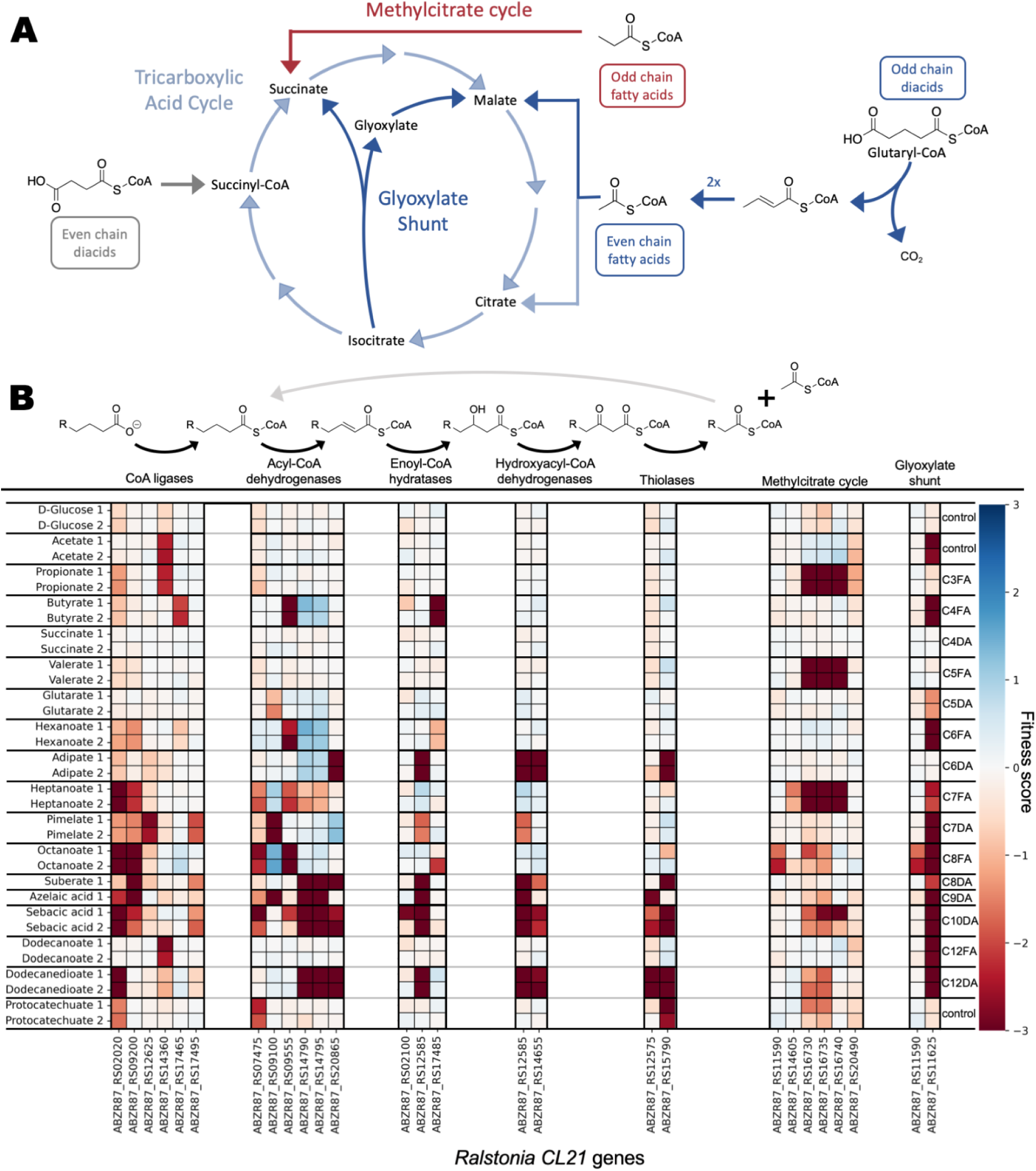
RB-TnSeq experiments identify genes involved in β-oxidation in *Ralstonia CL21.* (a) A diagram of the TCA cycle, the methylcitrate cycle, and the glyoxylate shunt. (b) One round of β-oxidation, with the genes predicted by Pfam to encode for β-oxidation enzymes, the glyoxylate shunt, and the methylcitrate cycle that had significant (|fitness| > 1 and |t-score| > 4) fitness phenotypes, and their corresponding fitness values on the conditions tested. Equivalent figures for *C. basilensis* 4G11, *P. bryophila* 376MFSha3.1, and *B. phytofirmans* PsJN can be found in Figures S7-9.

The first step of β-oxidation is activation with coenzyme A (CoA). From the data, we identified several acyl-CoA ligases involved in the metabolism of fatty and dicarboxylic acids. For example, ABZR87_RS12625 appeared to be a specific pimeloyl CoA-ligase, while ABZR87_RS09200 had a strong fitness defect on the C8 fatty and dicarboxylic acids, as well as the C9 dicarboxylic acid. Interestingly, ABZR87_RS02020 had a strong phenotype on the dicarboxylates dodecanedioic acid (C12-DA) and sebacic acid (C10-DA), but not on the C12 fatty acid dodecanoic acid. The library, like the wild-type strain, was unable to grow with the C10 fatty acid as a sole carbon source. However, ABZR87_RS02020 also had a strong fitness defect for the shorter chain length fatty acids octanoic and heptanoic acid. The CoA-ligase ABZR87_RS17495 seemed to have specificity for dicarboxylic acids, as it had significant phenotypes only for pimelate, suberate, azelaic acid, and sebacic acid carbon sources.

The second step of β-oxidation is catalyzed by an acyl-CoA dehydrogenase. Six genes predicted to code for this type of enzyme had significant fitness phenotypes, three of which had specific phenotypes for C8-C12 chain length dicarboxylic acids. Of those, ABZR87_RS14790 and ABZR87_RS14795, had the same fitness profile and may encode subunits of the same protein. The third, ABZR87_RS20865, also had a strong fitness phenotype on the C8-C12 dicarboxylic acids, in addition to a phenotype on the adipic acid condition. This could indicate that ABZR87_RS14795 and ABZR87_RS14790 act on the C8+ -CoAs, while ABZR87_RS20865 acts on their downstream product, adipyl-CoA. Likely for similar reasons, the predicted glutaryl-CoA acyl-CoA dehydrogenase ABZR87_RS09100 had a strong phenotype in the pimelic and azelaic acid conditions, however, its weaker phenotype in the glutaric acid condition was surprising and the subject of further examination. Another acyl-CoA dehydrogenase, ABZR87_RS09555, only had a significant phenotype with the fatty acid carbon source conditions.

The third and fourth steps of β-oxidation are hydration and dehydrogenation. The reaction is catalyzed by an enoyl-CoA hydratase and a 3-hydroxyacyl-CoA dehydrogenase, which can be separate enzymes or a single bifunctional one. Three genes predicted to code for enoyl-CoA hydratases had significant fitness phenotypes, and two genes were predicted to encode a hydroxyacyl-CoA dehydrogenase. Of these genes, ABZR87_RS14655 and ABZR87_RS12585 had strong fitness on the C6, C8-C10 and C12 dicarboxylic acids, while ABZR87_RS12585 also had significant fitness in the pimelic acid (C7-DA) condition. Only one had a significant fitness phenotype on fatty acids, ABZR87_RS17485, which appeared to be specific for the consumption of butyrate. No other predicted enoyl-CoA hydratases had strong phenotypes for the fatty acids, likely due to functional redundancies.

The final step in each round of β-oxidation is thiolysis, which yields acetyl-CoA and a fatty acyl-CoA that is two carbons shorter than the original compound. There were only two predicted thiolases in *R. CL21* with significant fitness phenotypes, ABZR87_RS15535 and ABZR87_RS12575. These genes both had significant fitness phenotypes on C8+ dicarboxylic acids, and ABZR87_RS15535 also had a significant fitness phenotype on the adipic acid and protocatechuic acid conditions. There were no thiolases with significant phenotypes with fatty acid carbon sources, again suggesting possible functional redundancies.

We also identified genes involved in the methylcitrate cycle and the glyoxylate shunt, two pathways that can be integral for efficient β-oxidation. The methylcitrate cycle processes propionyl-CoA, which is produced by the complete oxidation of odd-chain fatty acids. As expected, the predicted methylcitrate cycle genes only exhibited strong fitness defects with the odd-chain fatty acid carbon sources (Figure 4, Supplemental Table 11). The glyoxylate shunt bypasses the steps of the TCA cycle where CO_2_ is lost, enabling anabolism when the predominant carbon source is acetate/acetyl-CoA. The predicted glyoxylate shunt gene ABZR87_RS11625, a malate synthase, had strong phenotypes for the short chain fatty acids and all of the long-chain carbon sources (Figure 4). Even for the even chain dicarboxylates, which should result in succinyl-CoA that could feed directly into the tricarboxylic acid cycle, it appears that the glyoxylate shunt is necessary for optimal growth, likely due to the amount of acetyl-CoA released from the repeated rounds of β-oxidation to reach succinyl-CoA. This does not hold true for adipate, which only needs to undergo one round of β-oxidation to produce succinyl-CoA. There are multiple predicted redundancies (Supplemental Tables 11-12) for several genes in the methylcitrate cycle and glyoxylate shunt, which is the likely reason for many of those genes having few significant fitness phenotypes.

### Utilizing fitness data to improve short chain dicarboxylic acid consumption in R. CL21

The RB-TnSeq library for *R. CL21* exhibited growth with glutarate as the sole carbon source, which is unexpected considering that wild-type *R. CL21* exhibits no growth on glutarate (Figure 1, Figure 5C). This indicated that there existed mutants within the RB-TnSeq library that enabled utilization of glutarate.

**Figure 5:**
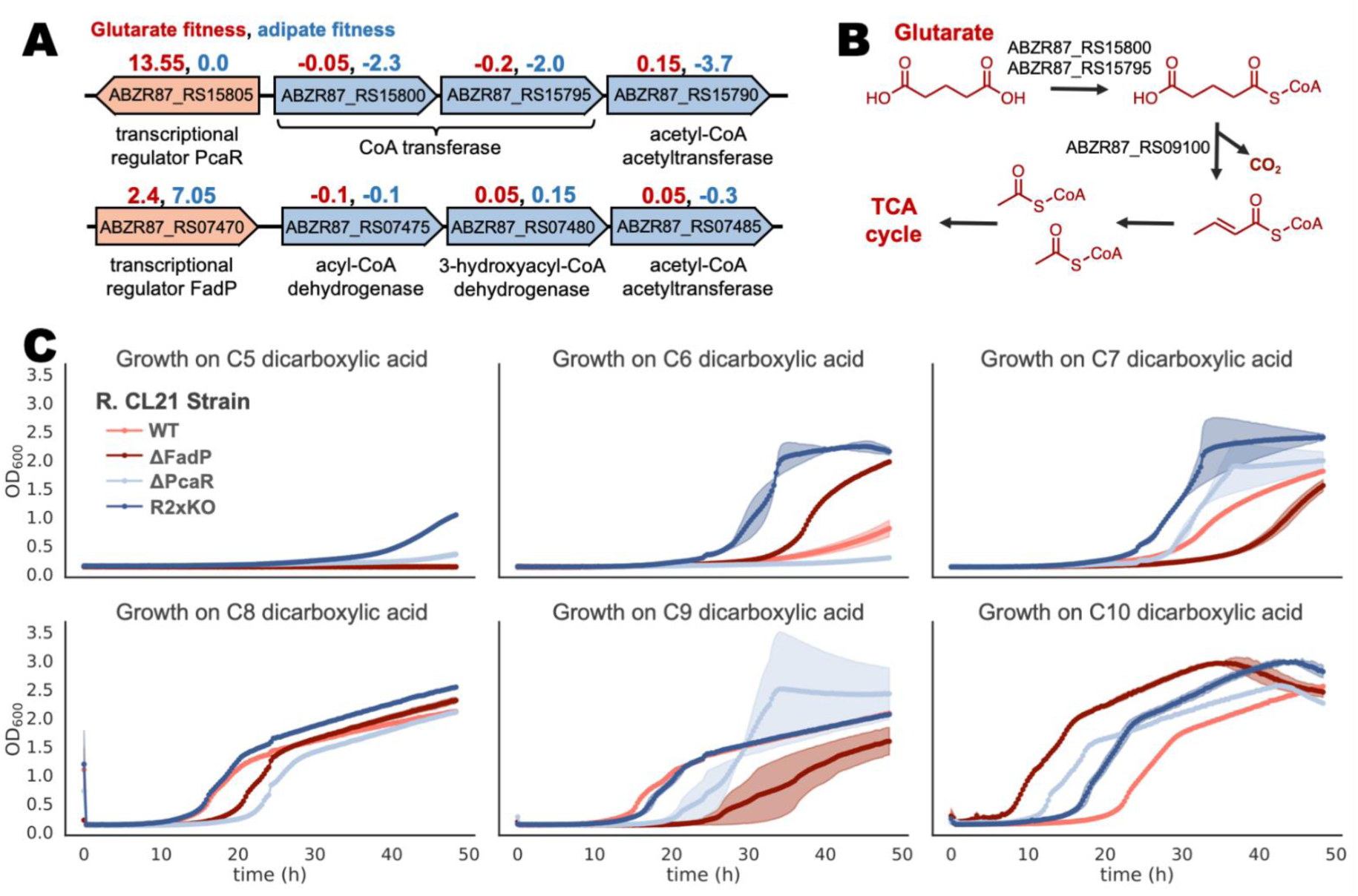
Deletion of two transcriptional regulators improves dicarboxylic acid consumption. A) Repressors with high positive fitness scores, ABZR87_RS15805 and ABZR87_RS07470, along with their predicted adjacent operons. Associated fitness values for glutarate (red, left) and adipate (blue, right) are shown. B) Proposed degradation pathway for glutarate. C) Growth curves for wild type, single, and double regulator knockout strains with dicarboxylic acids of varying chain lengths as sole carbon sources in MOPS-buffered minimal medium. The double knockout strain (R2xKO) grew on all chain lengths tested (n=3, error bars = 95% confidence interval).

Since transposon mutagenesis results in loss-of-function mutations, we examined the genes with positive fitness scores. An IclR-family transcriptional regulator, ABZR87_RS15805, had the highest fitness score with glutarate as sole carbon source, at 13.55, indicating that the average frequency of mutants with this gene disrupted by a barcoded transposon increased by 2^13.55^ times (Figure 5A). This family of transcription factors can be activators, repressors, or both (40), but in this case, the fitness data suggest that ABZR87_RS15805 likely functions as a repressor. Its neighboring operon, ABZR87_RS15775-15800, shares homology and synteny to the β-ketoadipate pathway of aromatic degradation, and the average fitness value across this operon with protocatechuate as a sole carbon source is strongly negative, at -3.9 (Figure 5A) (41, 42). However, the regulator itself has a strong positive score with protocatechuate as a carbon source, at +4.7. Since ABZR87_RS15805 and ABZR87_RS15775-15800 both have significant fitness scores on the same conditions, albeit opposite in magnitude, it is likely that ABZR87_RS15805 functions as the widely conserved β-ketoadipate pathway regulator PcaR, and represses the β-ketoadipate pathway ABZR87_RS15775-15800 in *R. CL21* (Figure 5A) (41).

Such a strong positive fitness score in the glutarate condition indicates that the mutants in ABZR87_RS15805 (*pcaR*) likely dominated the experiment population and may have skewed the data in such a way that explains the lack of strong fitness scores in predicted β-oxidation genes. For example, there is a significant fitness defect of -1.2 for the glutaryl-CoA dehydrogenase ABZR87_RS09100, but it had a much stronger score in the pimelic (-4.7) and azelaic (-4.6) acid experiments. These longer odd-chain dicarboxylic acids are β-oxidized to glutaryl-CoA, which then is dehydrogenated and decarboxylated by ABZR87_RS09100 and eventually converted to acetyl-CoA (Figure 4B).

Since it appears likely that *R. CL21* can metabolize glutaryl-CoA, it then follows that the inability to consume glutarate could be due to a lack of expression of a glutaryl-CoA ligase. The operon ABZR87_RS15775-15800 is likely constitutively expressed in *pcaR* mutants and contains ABZR87_RS15795-800, subunits of the enzyme predicted to transfer a CoA from succinate to β-keto-adipate (41). We hypothesized that ABZR87_RS15795-800 is more promiscuous than its regulator and can act on glutarate even though its regulator cannot detect it.

An in-frame deletion of *pcaR* did indeed enable growth with glutarate as a sole carbon source, albeit very little (Figure 5C). Additionally, when adipate was provided as the sole carbon source, the Δ*pcaR* mutant also displayed even less growth than that of the poorly growing wild-type strain. However, we identified a second transcriptional regulator, the TetR family repressor ABZR87_RS07470, with a positive fitness phenotype in the glutarate experiment (2.4) and a strong positive phenotype in the adipate experiment (7.05) (Figure 5A). Similar to PcaR, fitness data indicate that this transcription factor also represses an adjacent operon containing β-oxidation genes. ABZR87_RS07470 shares 95% identity and 92% coverage with the fatty-acid degradation repressor *fadP* identified in *Rastonia solanacearum* and proposed to regulate fatty-acid degradation in most species of the *Burkholderiales* order (43). Constructing a single deletion of ABZR87_RS07470 (*fadP*) resulted in improved growth with adipate as a carbon source but did not enable growth with glutarate (Figure 5C). However, a double knockout strain with in-frame deletions of both *pcaR* and *fadP* resulted in a strain that could grow robustly with all chain lengths of dicarboxylic acids tested, dubbed R2xKO (Figure 5C).

In *R. CL21*, both of these transcriptional repressors appear to be more specific than the genes that they regulate, masking cryptic metabolic capabilities. PcaR is known to respond to β-ketoadipate, and a homolog from *P. putida* (41.5% identity with 91% coverage to that of *R. CL21*) has been shown to respond to adipic acid, fumaric acid, glutaric acid, malic acid, and succinic acid (44). However, the very strong positive fitness phenotype of *pcaR* suggests that PcaR does not effectively de-repress its operon in the presence of glutarate in *R*. *CL21*. The fitness data does indicate that PcaR likely does respond to adipic acid in *R. CL21*. The CoA transferase ABZR87_RS15795-800 has negative fitness in the adipic acid condition, implying that it also acts on adipate, yet *pcaR* has a neutral phenotype (0.0) in the adipate condition, indicating that disrupting the repressor does not improve growth.

Similarly, the FadP inducer range might be more specific than the genes it regulates, however, this is would not be ascertainable from the fitness data alone, since FadP regulates multiple operons–many of which contain β-oxidation genes–throughout the genome in response to an unknown inducer. In *R. solanacearum* GMI1000, FadP regulates 12 operons composed of 27 genes, 15 of which encode for β-oxidation enzymes (43). We find that all of these genes were conserved in *R. CL21* with the most dissimilar protein still sharing 80% identity with 92% coverage to its *R. solanacearum* counterpart. It appears that regulation by FadP is conserved in *R. CL21* too. Taking the region 200 base pairs upstream of the operons in *R. CL21* and using MEME, we find a conserved motif that is highly similar to the reported binding site in *R. solanacearum* (43, 45). Eight of the twelve *R. CL21* operons have a significant fitness phenotype with at least one fatty or dicarboxylic acid condition, and one out of the four with insignificant phenotypes had no transposon insertions, likely because it was essential under the conditions the library was created in (Supplementary Table 13).

### Exploration of *R. CL21* as a potential host for plastics upcycling

To date, the bacterial strains used in upcycling dicarboxylic acids from plastics waste to value-added products required heterologously expressed β-oxidation, regulatory, and transport genes to enable dicarboxylic acid degradation (23). However, robust growth on dicarboxylic acids depends on more than just transport, β-oxidation, and their regulation (Figure 2A). Of the 501 genes with specific phenotypes solely on dicarboxylic acids, only 31.1% were predicted to be β-oxidation, regulatory, or transport genes. The remaining 68.9% of genes that were specifically necessary for robust growth on dicarboxylic acids may or may not be present in organisms engineered to consume dicarboxylic acids. These engineered hosts did not evolve to consume dicarboxylic acids, and therefore, their dicarboxylic acid metabolism is likely suboptimal. However, *R. CL21* naturally metabolizes multiple dicarboxylic acids, and the R2xKO strain is capable of robustly consuming all chain lengths of dicarboxylic acids. Therefore, this organism may be a promising host organism for upcycling dicarboxylic acids produced from plastics waste.

In contrast to most established host organisms, little research has been done with *R. CL21*. It was originally derived from the *Arabindopsis thaliana* root microbiome (46) and has been studied in the context of plant immune response to flagellar protein fragments (47). However, there is a lack of known tools or protocols available for its engineering. We sought to remedy this by characterizing several inducible and constitutive promoters in *R. CL21*, as well as determining what antibiotics could be used for alternative selections. We found that *R. CL21* was resistant to several antibiotics, with carbenicillin and apramycin failing to completely inhibit growth even at concentrations of 1000 mg/L. *R. CL21* was also naturally resistant to gentamicin at low to medium concentrations, therefore, gentamicin at 30 mg/L was routinely used to prevent contamination of *R. CL21* stock cultures. Kanamycin, chloramphenicol, and spectinomycin all demonstrated inhibition of *R. CL21* growth at concentrations >250 mg/L, and while only kanamycin 300 mg/L and the KanR resistance gene were used for selection in this work, the latter two antibiotics and their resistance markers would also be promising options (Figure S11A).

We also tested the strength of 13 constitutive promoters and 12 inducible systems from a previously published toolkit (48). These systems all control the expression of RFP from a kanamycin resistance backbone with a broad host-range origin (BBR1) and were introduced to *R. CL21* via electroporation. The constitutive promoters resulted in a broad range of expression levels, albeit, the promoters did behave differently in *R. CL21* compared to *E. coli* and *P. putida*, highlighting the importance of characterizing expression systems in the host organism of interest (Figure S11B) (48). Of the 13 inducible systems, six performed well in *R. CL21*: LacI/P_lacUV5_, XylS/P_M_, ChnR/P_ChnB_, Jungle Express, RhaR/P_Rha_, and AraC/P_BAD_.

After establishing tools to work with *R. CL21*, we next aimed to gain a more comprehensive understanding of the *R. CL21* wild-type (WT) and R2xKO strains’ abilities to utilize dicarboxylic acids, this time as mixtures instead of sole carbon sources. Using MOPS-buffered minimal medium as a base, we created two media. The first, referred to as ‘equimolar mix’, included 1 mM each of C4-C10, C12 dicarboxylic acids as the carbon source. The second medium, referred to as ‘mock mix’ consisted of the same dicarboxylic acids, but instead combined in a molar ratio that is more similar to that obtained from plastics waste, in this case, from chemical oxidation of high-density polyethylene (HDPE) (3) (Supplementary Table 14). We grew both *R. CL21* and R2xKO in these media, and collected samples after 12 and 24 hours to compare the depletion of dicarboxylic acids between the two different strains.

After 12 hours, there was no significant depletion of any of the dicarboxylic acids in the *R. CL21* WT experiments, and correspondingly, no noticeable growth of the cultures (Figures 6A, 6B). However, the R2xKO strain demonstrated appreciable degradation of succinic acid in both media, with no detectable succinic acid remaining in the equimolar mix medium after 12 hours. The R2xKO strain also noticeably increased in OD_600_ during the first 12 hours. After 24 hours, there were no detectable dicarboxylic acids in any of the R2xKO experiments. This was in contrast to the *R. CL21* WT strain, which mostly or entirely consumed the longer chain length dicarboxylic acids but had a significant amount of the C5-C7 dicarboxylic acids remaining after 24 hours of growth. There was some consumption of the glutaric acid by the WT strain after 24 hours, which contradicted our previous growth curves with glutarate as a sole source of carbon. However, with the mixed dicarboxylic acid carbon source provided in these experiments, some consumption of glutarate is expected, since, as previously discussed, the fitness data indicate that adipic acid likely induces the operon regulated by PcaR. The final OD_600_ of the WT strain was approximately half that of the R2xKO strain in both media (Figure 6C).

**Figure 6:**
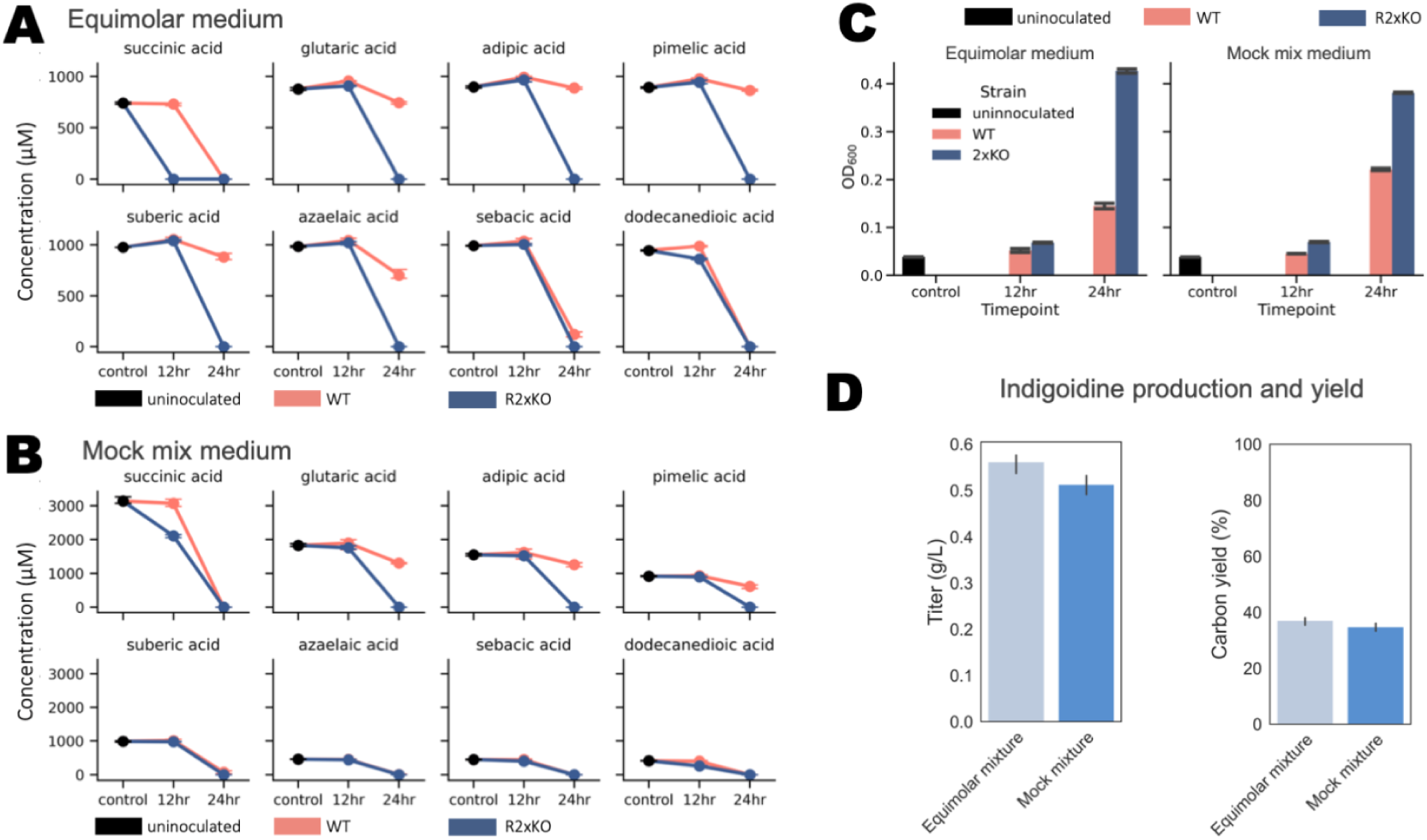
Dicarboxylic acid consumption and bioproduction of indigoidine, for all plots, n=3 and error bars = 95% confidence interval. A) Consumption of each dicarboxylic acid in an equimolar mixture of dicarboxylic acids by *R. CL21* WT and R2xKO over 24 hours, as measured by LC-MS. B) Consumption of each dicarboxylic acid by *R. CL21* WT and R2xKO over 24 hours, as measured by LC-MS. Initial concentrations of dicarboxylic acid were set to roughly what one could expect to obtain from the chemical oxidation of HDPE. C) Growth (as determined by OD_600_ measurements) of *R. CL21* WT and R2xKO over the course of the 24 hour consumption experiments. D) Plasmid-based production and yield of indigoidine in R2xKO from a mixture of dicarboxylic acids, measured colorimetrically (Figure S11A).

The robust and rapid utilization of dicarboxylic acids by the R2xKO strain, including the entirety of the mock mixture of dicarboxylic acids, indicates that this strain may have potential for plastics upcycling applications. To explore this further, we chose to engineer and test the bioproduction of indigoidine, a non-ribosomal peptide synthetase (NRPS)-derived blue pigment, from dicarboxylic acids as a sole source of carbon in our R2xKO strain (49). We selected indigoidine as our target compound for two reasons. First, indigoidine can be quantified by its absorbance at 612 nm, allowing us to rapidly measure the indigoidine titers (Figure S12) (50, 51). Second, its precursor, L-glutamine, is produced from α-ketoglutarate, an intermediate of the TCA cycle. Once converted from its apo-to its holo-form by the 4’-phosphopantetheinyl transferase Sfp, the NRPS BpsA condenses two molecules of L-glutamine into indigoidine (49). Due to the large flux through the TCA cycle during the β-oxidation of dicarboxylic acids, we expected that a target compound derived from a TCA cycle intermediate would be favorable for production from dicarboxylic acids.

After introducing a plasmid with BpsA and Sfp under the control of a medium-strength constitutive promoter to R2xKO, we grew the strain in equimolar and mock mix media with the selective marker kanamycin. The equimolar mixture contained a total of 1.35 g/L dicarboxylic acids and the mock mixture media contained 1.45 g/L dicarboxylic acids as their sole sources of carbon (Supplementary Table 14). After 36 hours, we extracted the indigoidine from the cultures and quantified it via absorbance at 612 nm. R2xKO produced 0.56 ± 0.02 g/L of indigoidine in the equimolar mix media and 0.51 ± 0.02 g/L in the mock mixture media (Figure 6D). This corresponds with 37% and 35% of the carbon present in the equimolar and mock mixture media being incorporated into indigoidine, respectively, and 62% and 60% of the estimated maximum theoretical yield from the provided carbon (Figure 6D). One possible explanation for the high titers could be the need to produce amino acids, which were not provided in the minimal media. Glutamine is the key component to nitrogen assimilation and the biosynthesis of amino acids, both acting as a key nitrogen donor to produce the amine group on amino acids and a direct precursor for several other amino acids (28). Since glutamine is necessary for the cell to survive, flux from the TCA cycle must be diverted into producing glutamine from α-ketoglutarate. It is possible that the consumption of glutamine by the indigoidine NRPS amplified this effect, forcing the microbe to direct more carbon into glutamine production in order to survive. This level of production, with minimal engineering, is very promising, and demonstrates that R2xKO could have potential as a host organism for plastics upcycling, especially for bioproducts derived from the TCA cycle.

## Conclusions

In four β-proteobacteria capable of consuming medium-to long-chain length dicarboxylic acids, RB-TnSeq identified genes involved in β-oxidation, transport, and regulation. The data indicated that oftentimes the genes that are required for metabolizing dicarboxylic acids are distinct from those required for metabolizing fatty acids. This dataset is more comprehensive than previous studies and will be useful for researchers aiming to engineer dicarboxylic acid production or catabolism in other organisms. In *R. CL21*, RB-TnSeq revealed cryptic metabolism for glutarate, which guided our construction of a deletion strain of two transcriptional regulators that enabled *R. CL21* to robustly consume short-chain dicarboxylic acids. A tool kit was established for *R. CL21* which should aid in future directions to explore *R. CL21* as a host for plastics upcycling.

As a proof of concept that *R. CL21* can be used as a host for plastics upcycling, we demonstrated the production of indigoidine from a mixture of dicarboxylic acids mimicking the ratios that can be obtained from plastics degradation. With only dicarboxylic acids as the source of carbon, we achieved titers of over 0.5 g/L and approximately 60% maximum theoretical yield from carbon. This demonstrated the efficiency of R2xKO at converting a TCA cycle intermediate into a bioproduct when dicarboxylic acids are the sole source of carbon. Future work could explore the production of other bioproducts derived from the TCA cycle, for instance, the biofuel isoprenol, which is produced from acetyl-CoA (52).

## Materials and Methods

### Media, chemicals and culture conditions

All cultures were grown at 30 °C with 200 rpm shaking, and grown either at a volume of 1 mL in 14 mL polypropylene round bottom tubes (Corning Falcon) or 3 mL in 55 mL borosilicate glass reusable culture tubes (VWR), unless otherwise stated. All overnight precultures were grown in lysogeny broth (LB) Miller medium (BD Biosciences). Modified MOPS-buffered minimal medium consisted of 32.5 μM CaCl_2_, 0.29 mM K_2_SO_4_, 1.32 mM K_2_HPO_4_, 8 μM FeCl_2_, 40 mM MOPS, 4 mM tricine, 0.01 mM FeSO_4_, 9.52 mM NH_4_Cl, 0.52 mM MgCl_2_, 50 mM NaCl, 0.03 μM (NH_4_)_6_Mo_7_O_24_, 4 μM H_3_BO_3_, 0.3 μM CoCl_2_, 0.1 μM CuSO_4_, 0.8 μM MnCl_2_, and 0.1 μM ZnSO_4_ (53). *R. CL21* cultures were supplemented with kanamycin (300 mg/L; Sigma-Aldrich) for selection or gentamicin (30 mg/L; Fisher Scientific) for maintenance. All other compounds were purchased through Sigma-Aldrich.

### Strains and plasmids

All the bacterial strains used in this study are listed in Table 1, and the plasmids used in this work are listed in Table 2. All strains and plasmids created in this work are available through the public instance of the JBEI registry (https://public-registry.jbei.org/folders/883). All plasmids were designed using Device Editor and Vector Editor software, while all primers used for the construction of plasmids were designed using j5 software (54, 55); (56). Plasmids were assembled via Gibson assembly using standard protocols (57). Plasmids were routinely isolated using a QIAprep Spin Miniprep kit (Qiagen), and all primers were purchased from Integrated DNA Technologies (IDT) (Coralville, IA).

**Table 1:** Strains used in this work.

| Strain | Description | JPUB ID | Reference |
| --- | --- | --- | --- |
| *Burkholderia phytofirmans* PsJN | Wild-type strain | - | (60) |
| *Cupriavidus basilensis* FW507-4G11 | Wild-type strain | - | (61) |
| *Dyella japonica* UNC79MFTsu3.2 | Wild-type strain | - | (46) |
| *Escherichia coli* BW25113 | Wild-type strain | - | (62) |
| *Herbaspirillum seropedicae* SmR1 | Wild-type strain | - | (63) |
| *Klebsiella michiganensis* M5a1 | Wild-type strain | - | (64) |
| *Paraburkholderia bryophila* 376MFSHa3.1 | Wild-type strain | - | (65) |
| *Pedobacter* sp. GW460-11-11-14-LB5 | Wild-type strain | - | (66) |
| *Pseudomonas simiae* WCS417 | Wild-type strain | - | (67) |
| *Sinorhizobium meliloti* 1021 | Wild-type strain | - | (68) |
| *Ralstonia* sp. UNC404CL21Col | Wild-type strain | - | (46, 47) |
| *R. CL21* ΔABZR87_RS15805, | Strain with complete internal in-frame deletion of ABZR87_RS15805 | JPUB_025979 | This work |
| *R. CL21* ΔABZR87_RS07470, | Strain with complete internal in-frame deletion of ABZR87_RS07470 | JPUB_025980 | This work |
| *R. CL21* ΔABZR87_RS15805 ΔABZR87_RS07470 (R2xKO) | Double knockout of ABZR87_RS15805 and ABZR87_RS07470 | JPUB_025981 | This work |
| *E. coli* XL1 Blue |  | - | Agilent |
| *E. coli* S17 |  | - | Agilent |

**Table 2:** Plasmids used in this work.

| Plasmid | Description | JPUB ID | Reference |
| --- | --- | --- | --- |
| pGingerBK-J23100 | Expression of RFP from J23100 promoter | - | (48) |
| pGingerBK-J23101 | Expression of RFP from J23101 promoter | - | (48) |
| pGingerBK-J23102 | Expression of RFP from J23102 promoter | - | (48) |
| pGingerBK-J23103 | Expression of RFP from J23103 promoter | - | (48) |
| pGingerBK-J23104 | Expression of RFP from J23104 promoter | - | (48) |
| pGingerBK-J23105 | Expression of RFP from J23105 promoter | - | (48) |
| pGingerBK-J23105 bpsA-sfp | Expression of bpsA-sfp from J23105 promoter | JPUB_026006 | This work |
| pGingerBK-J23107 | Expression of RFP from J23107 promoter | - | (48) |
| pGingerBK-J23109 | Expression of RFP from J23109 promoter | - | (48) |
| pGingerBK-J23110 | Expression of RFP from J23110 promoter | - | (48) |
| pGingerBK-J23111 | Expression of RFP from J23111 promoter | - | (48) |
| pGingerBK-J23113 | Expression of RFP from J23113 promoter | - | (48) |
| pGingerBK-J23114 | Expression of RFP from J23114 promoter | - | (48) |
| pGingerBK-J23119 | Expression of RFP from J23119 promoter | - | (48) |
| pGingerBK-LacUV5 | BBR1/Kanamycin backbone, IPTG-inducible | - | (48) |
| pGingerBK-XylS | BBR1/Kanamycin backbone,<br>Benzoate-inducible | - | (48) |
| pGingerBK-NahR | BBR1/Kanamycin resistance,<br>Salicylic acid-inducible | - | (48) |
| pGingerBK-JE | BBR1/Kanamycin resistance, Crystal violet-inducible | - | (48) |
| pGingerBK-RhaS | BBR1/Kanamycin resistance, Rhamnose-inducible | - | (48) |
| pMQk30 | Allelic exchange vector | - | (69) |
| pMQk30<br>ΔABZR87_RS15805 | pMQk30 derivative harboring 1-kb flanking regions of ABZR87_RS15805 | JPUB_025986 | This work |
| pMQk30<br>ΔABZR87_RS07470 | pMQk30 derivative harboring 1-kb flanking regions of ABZR87_RS07470 | JPUB_025984 | This work |

Construction of *R. CL21* deletion mutants via homologous recombination was performed following a protocol similar to that of *P. putida* (58). Allelic exchange vectors derived from the backbone pMQk30 (kanamycin resistance marker, *sacB* counterselection marker) were first introduced via conjugation. 2 mL of *E. coli* S17 harboring the allelic exchange vector pMQk30 (the donor strain) and *R. CL21* (the recipient strain) were washed twice with antibiotic-free LB via centrifugation for 3 minutes at 6,000 x g. Both the donor and the recipient strains were resuspended in 200 μL of LB. 100 μL of donor was combined with 50 μL of recipient, and MgSO_4_ was added at a concentration of 10 mM. The mixture was added to the center of a LB agar plate, with no spreading, and incubated at 30 °C for 24 hours. Next, the conjugation was scraped from the plate and resuspended in 1 mL of LB medium. 100 μL of this mixture was plated on an LB agar plate with kanamycin (300 mg/L) to select for the first crossover event in *R. CL21* and gentamicin (30 μg/mL) to counterselect for the *E. coli* donor strain. *R. CL21* colonies were picked from this plate and grown overnight in LB with gentamicin (30 μg/mL). These cultures were diluted 1:100 and 100 μL were plated on modified LB sucrose plates (per L, 100 g sucrose, 5 g yeast extract, 10 g tryptone, 15 g agar) to select for the crossing out of the backbone. Colonies were screened via colony pcr to distinguish between successful knockouts and wild-type revertants.

Plasmids were also introduced to *R. CL21* via electroporation, using a modified *Pseudomonas aeruginosa* protocol (59). For each transformation, 1 mL of cells grown for 24 hours was centrifuged for 3 min at 6000 x g, and the pellet was washed three times with 1 mL of room-temperature 300 mM sucrose. The pellet was then resuspended in 100 μL of 300 mM sucrose, and 50-200 ng of DNA was added prior to transferring to a 2 mm gap cuvette. Electroporation was performed with the BioRad MicroPulser at 2.5 kV.

### BarSeq assays

RB–TnSeq experiments utilized *R. CL21, B. phytofirmans PsJN, C. basilensis FW507-4G11, and P. bryophila 376MFSha3.1* libraries. The *B. phytofirmans PsJN, C. basilensis FW507-4G11, and P. bryophila 376MFSha3.1* libraries were described previously (32, 70), while the *R. CL21* RB-TnSeq library was constructed via conjugation using the barcoded mariner transposon vector PKMW3 (71). Libraries were thawed on ice, diluted in 25 mL of LB medium with kanamycin, and then grown to an OD_600_ of 0.5 at 30 °C, at which point three 1 mL aliquots were removed, pelleted, and stored at –80 °C as the time-zero control. The libraries were then washed once in MOPS minimal medium with no carbon source and then diluted 1:50 in MOPS minimal medium with 10 mM each carbon source tested. Cells were grown in 3 mL of medium in test tubes at 30 °C with shaking at 200 rpm. From 24 hours to 72 hours of growth, the cultures were inspected every 12 hours, and once cultures appeared visually turbid, the OD_600_ was recorded and a 500 μL aliquot was pelleted and stored at –80 °C until BarSeq analysis was performed as previously described (71, 72).

The fitness of each barcoded strain was determined by the log_2_ ratio of barcode reads in the sample to the barcode reads in the time zero control. The fitness score for a gene was the weighted average of these insertions in the central 10-90% of a gene. Fitness values are normalized such that the fitness value for a typical gene is zero. The primary statistical t-value is calculated from the estimated variance across different mutants of the same gene (71). Statistical t-values with absolute values greater than 4 were considered significant. All experiments were conducted in biological duplicate, and all fitness data are publicly available at http://fit.genomics.lbl.gov.

### Bioinformatic analyses

All statistical analyses were carried out using either the Python Scipy or Numpy libraries (73, 74). Biopython was also utilized in our analyses (75).To identify β-oxidation genes, HMMER version 3.3.2 (http://hmmer.org/) was used to scan the organism genomes against the relevant profile database files, shown in Supplementary Tables 6-12. The e-value cutoff was set to 1e-20. The database TransAAP was used to extract transport associated proteins from the BarSeq data set, and the set of Pfams used to identify regulators was inspired from the Pfams used in constructing the MiST database (Supplementary Table 3) (76, 77). PaperBlast and MetaCyc were utilized in the manual analysis of the data (78, 79).

Genes with significant fitness were defined as having a |fitness score| > 1 and |t-score| > 4 in at least one of the experimental conditions, while having a |fitness score| < 1 on the control condition (DL-lactate for *C. basilensis*, D-glucose for all other organisms).

Protein phylogenetic tree construction was completed by using MUSCLE with the super5 algorithm to align the protein sequences, followed by tree construction with the Phylo module of BioPython, using the UPGMA algorithm (80, 81). The UPGMA algorithm does not require an outgroup to be specified to root the tree, however, the *E. coli* protein sequence of AidB was added as a control.

Gene homology was calculated with BLAST+. First, a BLASTp database was created using makeblastdb containing all proteins with significant fitness from the four organisms. Then all sequences were compared to all others with the default E-value threshold (10) and a large maximum number of target sequences (1,000,000) to get a large number of pairwise comparisons. For plotting, an E-value threshold of 0.001 was used (82).

To compare fitness of multiple genes across organisms, we calculated pairwise the Pearson correlation coefficient between two genes’ fitness vectors. The fitness vector is constructed of the average fitness scores for the 20 growth conditions, with missing conditions masked. Correlation was calculated with the numpy ma.corrcoef function.

To address missing fitness scores in our dataset for t-SNE, which consists of vectors constructed from the average fitness scores of various organisms across 20 growth conditions, we used multiple imputation. This technique iteratively models each missing value as a function of other variables in the dataset, thus preserving the underlying statistical relationships. Specifically, we used KNNImputer from the scikit learn library with 2 nearest neighbors.

Following imputation, t-Distributed Stochastic Neighbor Embedding (t-SNE) was utilized to reduce the dimensionality of the imputed fitness vectors, facilitating a visual exploration of the data. This nonlinear technique is particularly adept at revealing patterns in multi-dimensional data by mapping high-dimensional data to a lower-dimensional space. Specifically, we used the TSNE module from the scikit learn library, with the perplexity set to 25, the number of iterations set to 50,000, and an early exaggeration of 100 (83). An interactive version of this analysis, with a display of gene name, annotation, and hyperlinks to the fitness browser can be accessed at [currently bokeh_tsne_subplots.html], and provides a useful way to visualize and browse the data across all four organisms.

### Plate-based growth assays

Both timepoint and kinetic assay were performed in plate format, and the BioTek Synergy H1 plate reader was used for all measurements.

For single timepoint OD and/or absorbance assays, overnight cultures were grown in LB, with the appropriate antibiotic when applicable. The cultures were then inoculated 1:100 into deep well plates. For the endpoint growth measurements of the different bacteria with fatty and dicarboxylic acid carbon sources, experiments were conducted in triplicate in deep 24-well plates with 2 mL of MOPS medium and 10 mM of each carbon source. Overnight cultures were washed twice in carbon-free MOPS minimal medium prior to inoculation. For the measurements of RFP expression from difference promoters in *R. CL21*, the experiments were conducted in deep 96-well plates with 0.5 mL of LB Kan300. The plates were sealed with a gas-permeable microplate-adhesive film (VWR). Optical density was measured by adding 100 uL of sample to a black sided, clear bottom 96-well plate, and measuring absorbance at 600 nm using a BioTek Synergy H1 plate reader. When applicable, RFP fluorescence was measured with an excitation wavelength of 535 nm, and emission wavelength of 620 nm, and a manually set gain of 100. For all continuous growth assays, overnight cultures were washed twice with MOPS minimal medium without any added carbon and diluted 1:100 into 500 μL of MOPS medium with 10 mM carbon source in 48-well plates (Falcon; 353072). The plates were sealed with BreathEasy transparent gas-permeable microplate-adhesive film (Diversified Biotech), and then, the optical density at 600 nm was monitored for 48 h in a BioTek Synergy H1 plate reader at 30 °C with fast continuous orbital shaking.

### Analytical determination of residual diacid content

Residual diacid content was analyzed by high-performance liquid chromatography mass spectrometry (HPLC-MS) on an Agilent 1260 Infinity II HPLC equipped with an Agilent MSD/XT single quadrupole detector. Samples were prepared by combining 250 μL of culture with an equal volume of ice-cold methanol, vortexing for 10 minutes, sonicating for 5 minutes, and centrifuging at 20,000 x g for 5 minutes. 100 μL of the supernatant was collected and stored at - 20 °C for later analysis. For this analysis, 2 μL was injected onto a Phenomenex Kinetex 2.6 μM XB-C18 100 Å column and separated with the following gradient with buffer A being composed of 0.1% formic acid in water and buffer B being composed of methanol with 0.1% formic acid:

| Time (min) | %B | Flow rate (mL/min) |
| --- | --- | --- |
| 0 | 5 | 0.425 |
| 6.5 | 97.1 | 0.650 |
| 7.8 | 97.1 | 0.650 |
| 8.2 | 5 | 0.650 |
| 12 | 5 | 0.650 |

The mass detector settings were as follows: 12.0 L/min drying gas flow, 35 psig nebulizer pressure, 350 °C drying gas temperature, and -3000 V capillary voltage. Diacids were detected with a negative mode scan from 80 to 250 m/z with a fragmentor voltage of 70, threshold abundance 150, step size 0.1 m/z, and speed of 193 u/sec.

Standards for each of the diacids with concentrations ranging from 5 to 2500 μM were analyzed as described above and used to construct a standard curve by measuring the area under the curve for each diacid’s extracted ion chromatogram (EIC) signal. The EIC values for samples were interpolated using these standard curves to determine the amount of residual diacid remaining in biological samples. The observed retention times for succinic acid, glutaric acid, adipic acid, pimelic acid, suberic acid, azelaic acid, sebacic acid, and dodecanedioic acid were 1.6 min, 2.3 min, 3.8 min, 4.8 min, 5.5 min, 6.0 min, 6.5 min, and 7.2 min, respectively.

### Production and quantification of indigoidine

An overnight culture in LB with Kan300 was washed twice with minimal medium and used to inoculate 3 mL production cultures in triplicate at a ratio of 1:100. To address residual indigoidine introduced via the inoculum, the concentration of indigoidine in the washed preculture was also determined and the appropriate value subtracted from the final titers. Cultures were harvested after 36 hours.

Indigoidine was extracted in DMSO and quantified via a colorimetric assay, as previously described (50, 51, 84). 3 mL of culture was collected and centrifuged for 2 minutes at 20,000 x g, and the supernatant subsequently removed. The resulting pellet was extracted in 1 mL of DMSO 3 times, via vigorous vortexing for 15 minutes, centrifugation for 2 minutes at 20,000 x g, and collection of the supernatant extract. The extract was combined, confirmed to a total volume of 3 mL, and used for the colorimetric assay.

A standard curve was prepared as previously described (50, 51). Calibration weights (Troemner, ASTM Class 4) were used to calibrate the scale and weigh three 20 mg aliquots of NMR-verified indigoidine (provided by the Mukhopadhyay group), which were then each combined with 40 mL of DMSO, wrapped in aluminum foil, and incubated overnight on a rocking platform to allow complete solubilization. A standard curve was generated in triplicate with 1.25-fold dilutions of this solution in DMSO. The absorbance of this standard curve at 612 nm, the maximally absorbed wavelength by indigoidine, was used to correlate OD_612_ to g/L of indigoidine, resulting in the formula: Y = 0.268x - 0.0109, which is similar to previously reported standard curve formulae for indigoidine (50, 51).

Absorbance measurements at 612 nm for 100 μL of the standard curve and samples were collected side-by-side in a clear 96-well flat-bottom plate (Corning Falcon) using a Biotek Synergy H1 plate reader. As a negative control, extract of wild-type *R. CL21* via the same procedure resulted in the same OD_612_ values as DMSO alone, indicating that any residual cell mass did not contribute to OD_612_. Full absorbance spectra of the extracted sample and the standard were collected, and showed no significant differences (Supplementary Figure 11).

### Flux Balance Estimate of Max Theoretical Yield

We used flux balance analysis to estimate maximum theoretical yield of indigoidine from dicarboxylic acids. This analysis was done in Python with CobraPy version 0.29.0 (85). We started with an *E. coli* central carbon metabolism model which includes a well conserved TCA cycle, BiGG model iJO1366 (86, 87). We further modified the model by knocking out ATP maintenance cost, adding indigoidine synthesis, and adding dicarboxylic acid transport and catabolism to TCA cycle intermediates.

For each diacid length (C5 and larger) we added reactions for transport, CoA ligation, and β-oxidation down to a smaller acyl-CoA. For dicarboxylic acid transport, we used the fitness data to ascertain which transporter types were responsible for each chain length of dicarboxylic acid (Figure S3A). These data, combined with gene homology to known transporters, indicated that cation symporters were responsible for the import of C4-C6 dicarboxylic acids which we modeled as 2-proton symporters, and ABC transporters were responsible for the import of C7+ dicarboxylic acids which we modeled as 2-ATP dependent (88). We modeled the β-oxidation of longer chain dicarboxylic acids down to one of two products – the C5 dicarboxylic acid glutarate for odd-chain length dicarboxylic acids and the C4 dicarboxylic acid succinyl-CoA for the even chain length dicarboxylic acids. Succinyl-CoA is an intermediate of the TCA cycle, and could be directly incorporated into the original model. For glutaryl-CoA, we included the reactions for the decarboxylative glutaryl-CoA degradation pathway from the *P. putida* model iJN1463 (89). We choose to only include the decarboxylative glutaryl-CoA degradation pathway because there were *R. CL21* genes homologous to the *P. putida* genes for this pathway with significant fitness phenotypes, but no *R. CL21* genes homologous to those of the CoA-independent pathway in *P. putida* (58).

We used our modified model, which can be found in Supplemental File 4, to estimate the maximum theoretical yield for each chain length of dicarboxylic acid. We then used the weighted averages of these values to find the estimated maximum theoretical yield for each medium’s mixture of dicarboxylic acid carbon sources.

## Acknowledgements

We thank Morgan Price for his assistance with RB-TnSeq analysis, Leah Keiser for her advice on LC/MS analysis, and Dr. Alberto Nava, Dr. Peter Winegar, Maria Astolfi, and Lucas Waldburger for their advice and assistance. Dr. Mitchell Thompson is a Simons Foundation Awardee of the Life Sciences Research Foundation. This work was part of the DOE Joint BioEnergy Institute (https://www.jbei.org) supported by the U.S. Department of Energy, Office of Science, Office of Biological and Environmental Research, supported by the U.S. Department of Energy, Energy Efficiency and Renewable Energy, Bioenergy Technologies Office, through contract DE-AC02-05CH11231 between Lawrence Berkeley National Laboratory and the U.S. Department of Energy. The views and opinions of the authors expressed herein do not necessarily state or reflect those of the United States Government or any agency thereof. Neither the United States Government nor any agency thereof, nor any of their employees, makes any warranty, expressed or implied, or assumes any legal liability or responsibility for the accuracy, completeness, or usefulness of any information, apparatus, product, or process disclosed, or represents that its use would not infringe privately owned rights.

Conceptualization: A.N.P. and M.G.T.; methodology: A.N.P., G.A.H. J.B.R. M.G.T., A.M.D.; investigation: A.N.P., J. M. L., C.N.H., G.A.H., A.V., J.B.R., J.M., M.R.I, M.S., M.G.T., A.M.D.; writing – original draft: A.N.P. and J.M.L.; writing – review and editing: all authors; resources and supervision: M.G.T., P.M.S., J.D.K.

J.D.K. has financial interests in Ansa Biotechnologies, Apertor Pharma, Berkeley Yeast, Demetrix, Lygos, Napigen, ResVita Bio, and Zero Acre Farms.

## Bibliography

1. Chae TU, Ahn JH, Ko Y-S, Kim JW, Lee JA, Lee EH, Lee SY. 2020. Metabolic engineering for the production of dicarboxylic acids and diamines. Metab Eng 58:2–16.

2. Yu J-L, Qian Z-G, Zhong J-J. 2018. Advances in bio-based production of dicarboxylic acids longer than C4. Eng Life Sci 18:668–681.

3. Sullivan KP, Werner AZ, Ramirez KJ, Ellis LD, Bussard JR, Black BA, Brandner DG, Bratti F, Buss BL, Dong X, Haugen SJ, Ingraham MA, Konev MO, Michener WE, Miscall J, Pardo I, Woodworth SP, Guss AM, Román-Leshkov Y, Stahl SS, Beckham GT. 2022. Mixed plastics waste valorization through tandem chemical oxidation and biological funneling. Science 378:207–211.

4. Pinsuwan K, Opaprakasit P, Petchsuk A, Dubas L, Opaprakasit M. 2023. Chemical recycling of high-density polyethylene (HDPE) wastes by oxidative degradation to dicarboxylic acids and their use as value-added curing agents for acrylate-based materials. Polym Degrad Stab 210:110306.

5. Bäckström E, Odelius K, Hakkarainen M. 2017. Trash to Treasure: Microwave-Assisted Conversion of Polyethylene to Functional Chemicals. Ind Eng Chem Res 56:14814–14821.

6. Wang K, Jia R, Cheng P, Shi L, Wang X, Huang L. 2023. Highly Selective Catalytic Oxi-upcycling of Polyethylene to Aliphatic Dicarboxylic Acid under a Mild Hydrogen-Free Process. Angew Chem 135.

7. Geyer R, Jambeck JR, Law KL. 2017. Production, use, and fate of all plastics ever made. Sci Adv 3:e1700782.

8. Tiso T, Winter B, Wei R, Hee J, de Witt J, Wierckx N, Quicker P, Bornscheuer UT, Bardow A, Nogales J, Blank LM. 2022. The metabolic potential of plastics as biotechnological carbon sources - Review and targets for the future. Metab Eng 71:77–98.

9. Lee S, Lee YR, Kim SJ, Lee J-S, Min K. 2023. Recent advances and challenges in the biotechnological upcycling of plastic wastes for constructing a circular bioeconomy. Chemical Engineering Journal 454:140470.

10. Gu S, Zhu F, Zhang L, Wen J. 2024. Mid-Long Chain Dicarboxylic Acid Production via Systems Metabolic Engineering: Progress and Prospects. J Agric Food Chem 72:5555– 5573.

11. Li W, Shen X, Wang J, Sun X, Yuan Q. 2021. Engineering microorganisms for the biosynthesis of dicarboxylic acids. Biotechnol Adv 48:107710.

12. Li G, Huang D, Sui X, Li S, Huang B, Zhang X, Wu H, Deng Y. 2020. Advances in microbial production of medium-chain dicarboxylic acids for nylon materials. React Chem Eng 5:221–238.

13. Song J-W, Lee J-H, Bornscheuer UT, Park J-B. 2014. Microbial Synthesis of Medium-Chain α,ω-Dicarboxylic Acids and ω-Aminocarboxylic Acids from Renewable Long-Chain Fatty Acids. Adv Synth Catal 356:1782–1788.

14. Haushalter RW, Phelan RM, Hoh KM, Su C, Wang G, Baidoo EEK, Keasling JD. 2017. Production of Odd-Carbon Dicarboxylic Acids in Escherichia coli Using an Engineered Biotin-Fatty Acid Biosynthetic Pathway. J Am Chem Soc 139:4615–4618.

15. Hagen A, Poust S, Rond T de, Fortman JL, Katz L, Petzold CJ, Keasling JD. 2016. Engineering a polyketide synthase for in vitro production of adipic acid. ACS Synth Biol 5:21–27.

16. Kim S, Gonzalez R. 2018. Selective production of decanoic acid from iterative reversal of β-oxidation pathway. Biotechnol Bioeng 115:1311–1320.

17. Zhao M, Huang D, Zhang X, Koffas MAG, Zhou J, Deng Y. 2018. Metabolic engineering of Escherichia coli for producing adipic acid through the reverse adipate-degradation pathway. Metab Eng 47:254–262.

18. Bao T, Qian Y, Xin Y, Collins JJ, Lu T. 2023. Engineering microbial division of labor for plastic upcycling. Nat Commun 14:5712.

19. Yoshioka T, Sato T, Okuwaki A. 1994. Hydrolysis of waste PET by sulfuric acid at 150°C for a chemical recycling. J Appl Polym Sci 52:1353–1355.

20. Tournier V, Topham CM, Gilles A, David B, Folgoas C, Moya-Leclair E, Kamionka E, Desrousseaux ML, Texier H, Gavalda S, Cot M, Guémard E, Dalibey M, Nomme J, Cioci G, Barbe S, Chateau M, André I, Duquesne S, Marty A. 2020. An engineered PET depolymerase to break down and recycle plastic bottles. Nature 580:216–219.

21. Valenzuela-Ortega M, Suitor JT, White MFM, Hinchcliffe T, Wallace S. 2023. Microbial upcycling of waste PET to adipic acid. ACS Cent Sci 9:2057–2063.

22. Brandenberg OF, Schubert OT, Kruglyak L. 2022. Towards synthetic PETtrophy: Engineering Pseudomonas putida for concurrent polyethylene terephthalate (PET) monomer metabolism and PET hydrolase expression. Microb Cell Fact 21:119.

23. Ackermann YS, Li W-J, Op de Hipt L, Niehoff P-J, Casey W, Polen T, Köbbing S, Ballerstedt H, Wynands B, O’Connor K, Blank LM, Wierckx N. 2021. Engineering adipic acid metabolism in Pseudomonas putida. Metab Eng 67:29–40.

24. Ackermann YS, de Witt J, Mezzina MP, Schroth C, Polen T, Nikel PI, Wynands B, Wierckx N. 2024. Bio-upcycling of even and uneven medium-chain-length diols and dicarboxylates to polyhydroxyalkanoates using engineered Pseudomonas putida. Microb Cell Fact 23:54.

25. Parke D, Garcia MA, Ornston LN. 2001. Cloning and genetic characterization of dca genes required for beta-oxidation of straight-chain dicarboxylic acids in Acinetobacter sp. strain ADP1. Appl Environ Microbiol 67:4817–4827.

26. Strittmatter CS, Eggers J, Biesgen V, Hengsbach J-N, Sakatoku A, Albrecht D, Riedel K, Steinbüchel A. 2022. Insights into the Degradation of Medium-Chain-Length Dicarboxylic Acids in Cupriavidus necator H16 Reveal β-Oxidation Differences between Dicarboxylic Acids and Fatty Acids. Appl Environ Microbiol 88:e0187321.

27. Harrison FH, Harwood CS. 2005. The pimFABCDE operon from Rhodopseudomonas palustris mediates dicarboxylic acid degradation and participates in anaerobic benzoate degradation. Microbiology (Reading, Engl) 151:727–736.

28. Schmidt M, Pearson AN, Incha MR, Thompson MG, Baidoo EEK, Kakumanu R, Mukhopadhyay A, Shih PM, Deutschbauer AM, Blank LM, Keasling JD. 2022. Nitrogen Metabolism in Pseudomonas putida: Functional Analysis Using Random Barcode Transposon Sequencing. Appl Environ Microbiol 88:e0243021.

29. Thompson MG, Pearson AN, Barajas JF, Cruz-Morales P, Sedaghatian N, Costello Z, Garber ME, Incha MR, Valencia LE, Baidoo EEK, Martin HG, Mukhopadhyay A, Keasling JD. 2020. Identification, Characterization, and Application of a Highly Sensitive Lactam Biosensor from Pseudomonas putida. ACS Synth Biol 9:53–62.

30. Thompson MG, Incha MR, Pearson AN, Schmidt M, Sharpless WA, Eiben CB, Cruz-Morales P, Blake-Hedges JM, Liu Y, Adams CA, Haushalter RW, Krishna RN, Lichtner P, Blank LM, Mukhopadhyay A, Deutschbauer AM, Shih PM, Keasling JD. 2020. Fatty acid and alcohol metabolism in Pseudomonas putida: functional analysis using random barcode transposon sequencing. Appl Environ Microbiol 86:e01665–20.

31. Harwood CS, Parales RE. 1996. The beta-ketoadipate pathway and the biology of self-identity. Annu Rev Microbiol 50:553–590.

32. Price MN, Wetmore KM, Waters RJ, Callaghan M, Ray J, Liu H, Kuehl JV, Melnyk RA, Lamson JS, Suh Y, Carlson HK, Esquivel Z, Sadeeshkumar H, Chakraborty R, Zane GM, Rubin BE, Wall JD, Visel A, Bristow J, Blow MJ, Deutschbauer AM. 2018. Mutant phenotypes for thousands of bacterial genes of unknown function. Nature 557:503–509.

33. Swigonová Z, Mohsen A-W, Vockley J. 2009. Acyl-CoA dehydrogenases: Dynamic history of protein family evolution. J Mol Evol 69:176–193.

34. Rippa V, Duilio A, di Pasquale P, Amoresano A, Landini P, Volkert MR. 2011. Preferential DNA damage prevention by the E. coli AidB gene: A new mechanism for the protection of specific genes. DNA Repair (Amst) 10:934–941.

35. Voskuil MI. 2013. Mycobacterium tuberculosis cholesterol catabolism requires a new class of acyl coenzyme A dehydrogenase. J Bacteriol 195:4319–4321.

36. Wipperman MF, Yang M, Thomas ST, Sampson NS. 2013. Shrinking the FadE proteome of Mycobacterium tuberculosis: insights into cholesterol metabolism through identification of an α2β2 heterotetrameric acyl coenzyme A dehydrogenase family. J Bacteriol 195:4331–4341.

37. Thomas ST, Sampson NS. 2013. Mycobacterium tuberculosis utilizes a unique heterotetrameric structure for dehydrogenation of the cholesterol side chain. Biochemistry 52:2895–2904.

38. Thompson MG, Blake-Hedges JM, Cruz-Morales P, Barajas JF, Curran SC, Harris NC, Benites VT, Gin JW, Eiben CB, Sharpless WA, Krishna RN, Baidoo E, Petzold CJ, Arkin AP, Deutschbauer AM, Keasling JD. 2018. Massively parallel fitness profiling reveals multiple novel enzymes in Pseudomonas putida lysine metabolism. BioRxiv 10.1101/450254.

39. Storz G, Tartaglia LA. 1992. OxyR: a regulator of antioxidant genes. J Nutr 122:627–630.

40. Krell T, Molina-Henares AJ, Ramos JL. 2006. The IclR family of transcriptional activators and repressors can be defined by a single profile. Protein Sci 15:1207–1213.

41. Parales RE, Harwood CS. 1993. Regulation of the pcaIJ genes for aromatic acid degradation in Pseudomonas putida. J Bacteriol 175:5829–5838.

42. Jiménez JI, Miñambres B, García JL, Díaz E. 2002. Genomic analysis of the aromatic catabolic pathways from Pseudomonas putida KT2440. Environ Microbiol 4:824–841.

43. Kazakov AE, Rodionov DA, Alm E, Arkin AP, Dubchak I, Gelfand MS. 2009. Comparative genomics of regulation of fatty acid and branched-chain amino acid utilization in proteobacteria. J Bacteriol 191:52–64.

44. Pham C, Nasr MA, Skarina T, Di Leo R, Kwan DH, Martin VJJ, Stogios PJ, Mahadevan R, Savchenko A. 2024. Functional and structural characterization of an IclR family transcription factor for the development of dicarboxylic acid biosensors. FEBS J 10.1111/febs.17149.

45. Bailey TL, Elkan C. 1994. Fitting a mixture model by expectation maximization to discover motifs in biopolymers. Proc Int Conf Intell Syst Mol Biol 2:28–36.

46. Levy A, Salas Gonzalez I, Mittelviefhaus M, Clingenpeel S, Herrera Paredes S, Miao J, Wang K, Devescovi G, Stillman K, Monteiro F, Rangel Alvarez B, Lundberg DS, Lu T-Y, Lebeis S, Jin Z, McDonald M, Klein AP, Feltcher ME, Rio TG, Grant SR, Dangl JL. 2017. Genomic features of bacterial adaptation to plants. Nat Genet 50:138–150.

47. Colaianni NR, Parys K, Lee H-S, Conway JM, Kim NH, Edelbacher N, Mucyn TS, Madalinski M, Law TF, Jones CD, Belkhadir Y, Dangl JL. 2021. A complex immune response to flagellin epitope variation in commensal communities. Cell Host Microbe 29:635–649.e9.

48. Pearson AN, Thompson MG, Kirkpatrick LD, Ho C, Vuu KM, Waldburger LM, Keasling JD, Shih PM. 2023. The pGinger Family of Expression Plasmids. Microbiol Spectr 11:e0037323.

49. Wehrs M, Gladden JM, Liu Y, Platz L, Prahl J-P, Moon J, Papa G, Sundstrom E, Geiselman GM, Tanjore D, Keasling JD, Pray TR, Simmons BA, Mukhopadhyay A. 2019. Sustainable bioproduction of the blue pigment indigoidine: Expanding the range of heterologous products in *R. toruloides* to include non-ribosomal peptides. Green Chem 21:3394–3406.

50. Banerjee D, Eng T, Lau AK, Sasaki Y, Wang B, Chen Y, Prahl J-P, Singan VR, Herbert RA, Liu Y, Tanjore D, Petzold CJ, Keasling JD, Mukhopadhyay A. 2020. Genome-scale metabolic rewiring improves titers rates and yields of the non-native product indigoidine at scale. Nat Commun 11:5385.

51. Eng T, Banerjee D, Menasalvas J, Chen Y, Gin J, Choudhary H, Baidoo E, Chen JH, Ekman A, Kakumanu R, Diercks YL, Codik A, Larabell C, Gladden J, Simmons BA, Keasling JD, Petzold CJ, Mukhopadhyay A. 2023. Maximizing microbial bioproduction from sustainable carbon sources using iterative systems engineering. Cell Rep 42:113087.

52. Chou HH, Keasling JD. 2012. Synthetic pathway for production of five-carbon alcohols from isopentenyl diphosphate. Appl Environ Microbiol 78:7849–7855.

53. LaBauve AE, Wargo MJ. 2012. Growth and laboratory maintenance of Pseudomonas aeruginosa. Curr Protoc Microbiol Chapter 6:Unit 6E.1.

54. Chen J, Densmore D, Ham TS, Keasling JD, Hillson NJ. 2012. DeviceEditor visual biological CAD canvas. J Biol Eng 6:1.

55. Ham TS, Dmytriv Z, Plahar H, Chen J, Hillson NJ, Keasling JD. 2012. Design, implementation and practice of JBEI-ICE: an open source biological part registry platform and tools. Nucleic Acids Res 40:e141.

56. Hillson NJ, Rosengarten RD, Keasling JD. 2012. j5 DNA assembly design automation software. ACS Synth Biol 1:14–21.

57. Gibson DG, Young L, Chuang R-Y, Venter JC, Hutchison CA, Smith HO. 2009. Enzymatic assembly of DNA molecules up to several hundred kilobases. Nat Methods 6:343–345.

58. Thompson MG, Blake-Hedges JM, Cruz-Morales P, Barajas JF, Curran SC, Eiben CB, Harris NC, Benites VT, Gin JW, Sharpless WA, Twigg FF, Skyrud W, Krishna RN, Pereira JH, Baidoo EEK, Petzold CJ, Adams PD, Arkin AP, Deutschbauer AM, Keasling JD. 2019. Massively Parallel Fitness Profiling Reveals Multiple Novel Enzymes in Pseudomonas putida Lysine Metabolism. MBio 10.

59. Choi K-H, Kumar A, Schweizer HP. 2006. A 10-min method for preparation of highly electrocompetent *Pseudomonas aeruginosa* cells: application for DNA fragment transfer between chromosomes and plasmid transformation. J Microbiol Methods 64:391–397.

60. Weilharter A, Mitter B, Shin MV, Chain PSG, Nowak J, Sessitsch A. 2011. Complete genome sequence of the plant growth-promoting endophyte Burkholderia phytofirmans strain PsJN. J Bacteriol 193:3383–3384.

61. Ray J, Waters RJ, Skerker JM, Kuehl JV, Price MN, Huang J, Chakraborty R, Arkin AP, Deutschbauer A. 2015. Complete Genome Sequence of Cupriavidus basilensis 4G11, Isolated from the Oak Ridge Field Research Center Site. Genome Announc 3.

62. Grenier F, Matteau D, Baby V, Rodrigue S. 2014. Complete Genome Sequence of Escherichia coli BW25113. Genome Announc 2.

63. Pedrosa FO, Monteiro RA, Wassem R, Cruz LM, Ayub RA, Colauto NB, Fernandez MA, Fungaro MHP, Grisard EC, Hungria M, Madeira HMF, Nodari RO, Osaku CA, Petzl-Erler ML, Terenzi H, Vieira LGE, Steffens MBR, Weiss VA, Pereira LFP, Almeida MIM, Souza EM. 2011. Genome of Herbaspirillum seropedicae strain SmR1, a specialized diazotrophic endophyte of tropical grasses. PLoS Genet 7:e1002064.

64. Yu Z, Li S, Li Y, Jiang Z, Zhou J, An Q. 2018. Complete genome sequence of N2-fixing model strain Klebsiella sp. nov. M5al, which produces plant cell wall-degrading enzymes and siderophores. Biotechnol Rep (Amst) 17:6–9.

65. Nethery MA, Hidalgo-Cantabrana C, Roberts A, Barrangou R. 2022. CRISPR-based engineering of phages for in situ bacterial base editing. Proc Natl Acad Sci USA 119:e2206744119.

66. Price MN, Deutschbauer AM, Arkin AP. 2022. Filling gaps in bacterial catabolic pathways with computation and high-throughput genetics. PLoS Genet 18:e1010156.

67. Vela AI, Gutiérrez MC, Falsen E, Rollán E, Simarro I, García P, Domínguez L, Ventosa A, Fernández-Garayzábal JF. 2006. Pseudomonas simiae sp. nov., isolated from clinical specimens from monkeys (Callithrix geoffroyi). Int J Syst Evol Microbiol 56:2671–2676.

68. Entcheva P, Phillips DA, Streit WR. 2002. Functional analysis of Sinorhizobium meliloti genes involved in biotin synthesis and transport. Appl Environ Microbiol 68:2843–2848.

69. Thompson M, Kirkpatrick LD, Geiselman GM, Waldburger LM, Pearson AN, Szarzanowicz M, Vuu KM, Markel K, Hummel NF, Suazo DD, Tahmin C, Cui R, Liu S, Cevallos J, Pannu H, Liu D, Gin JW, Chen Y, Petzold CJ, Gladden J, Shih PM. 2023. Genetically refactored Agrobacterium-mediated transformation. BioRxiv 10.1101/2023.10.13.561914.

70. Price MN, Ray J, Iavarone AT, Carlson HK, Ryan EM, Malmstrom RR, Arkin AP, Deutschbauer AM. 2019. Oxidative pathways of deoxyribose and deoxyribonate catabolism. mSystems 4.

71. Wetmore KM, Price MN, Waters RJ, Lamson JS, He J, Hoover CA, Blow MJ, Bristow J, Butland G, Arkin AP, Deutschbauer A. 2015. Rapid quantification of mutant fitness in diverse bacteria by sequencing randomly bar-coded transposons. MBio 6:e00306–15.

72. Rand JM, Pisithkul T, Clark RL, Thiede JM, Mehrer CR, Agnew DE, Campbell CE, Markley AL, Price MN, Ray J, Wetmore KM, Suh Y, Arkin AP, Deutschbauer AM, Amador-Noguez D, Pfleger BF. 2017. A metabolic pathway for catabolizing levulinic acid in bacteria. Nat Microbiol 2:1624–1634.

73. Virtanen P, Gommers R, Oliphant TE, Haberland M, Reddy T, Cournapeau D, Burovski E, Peterson P, Weckesser W, Bright J, van der Walt SJ, Brett M, Wilson J, Millman KJ, Mayorov N, Nelson ARJ, Jones E, Kern R, Larson E, Carey CJ, SciPy 1.0 Contributors. 2020. SciPy 1.0: fundamental algorithms for scientific computing in Python. Nat Methods 17:261–272.

74. Harris CR, Millman KJ, van der Walt SJ, Gommers R, Virtanen P, Cournapeau D, Wieser E, Taylor J, Berg S, Smith NJ, Kern R, Picus M, Hoyer S, van Kerkwijk MH, Brett M, Haldane A, Del Río JF, Wiebe M, Peterson P, Gérard-Marchant P, Oliphant TE. 2020. Array programming with NumPy. Nature 585:357–362.

75. Cock PJA, Antao T, Chang JT, Chapman BA, Cox CJ, Dalke A, Friedberg I, Hamelryck T, Kauff F, Wilczynski B, de Hoon MJL. 2009. Biopython: freely available Python tools for computational molecular biology and bioinformatics. Bioinformatics 25:1422–1423.

76. Gumerov VM, Ortega DR, Adebali O, Ulrich LE, Zhulin IB. 2020. MiST 3.0: an updated microbial signal transduction database with an emphasis on chemosensory systems. Nucleic Acids Res 48:D459–D464.

77. Elbourne LDH, Tetu SG, Hassan KA, Paulsen IT. 2017. TransportDB 2.0: a database for exploring membrane transporters in sequenced genomes from all domains of life. Nucleic Acids Res 45:D320–D324.

78. Price MN, Arkin AP. 2017. PaperBLAST: Text Mining Papers for Information about Homologs. mSystems 2.

79. Caspi R, Billington R, Keseler IM, Kothari A, Krummenacker M, Midford PE, Ong WK, Paley S, Subhraveti P, Karp PD. 2020. The MetaCyc database of metabolic pathways and enzymes - a 2019 update. Nucleic Acids Res 48:D445–D453.

80. Edgar RC. 2004. MUSCLE: multiple sequence alignment with high accuracy and high throughput. Nucleic Acids Res 32:1792–1797.

81. Talevich E, Invergo BM, Cock PJA, Chapman BA. 2012. Bio.Phylo: a unified toolkit for processing, analyzing and visualizing phylogenetic trees in Biopython. BMC Bioinformatics 13:209.

82. Stormo GD. 2009. An introduction to sequence similarity (“homology”) searching. Curr Protoc Bioinformatics Chapter 3:Unit 3.1 3.1.1-7.

83. Pedregosa F, Varoquaux G, Gramfort A, Michel V, Thirion B, Grisel O, Blondel M, Prettenhofer P, Weiss R, Dubourg V, Vanderplas J, Passos A, Cournapeau D, Brucher M, Perrot M, Duchesnay E. 2011. Scikit-learn: Machine Learning in Python. J Mach Learn Res 12:2825–2830.

84. Yu D, Xu F, Valiente J, Wang S, Zhan J. 2013. An indigoidine biosynthetic gene cluster from Streptomyces chromofuscus ATCC 49982 contains an unusual IndB homologue. J Ind Microbiol Biotechnol 40:159–168.

85. Ebrahim A, Lerman JA, Palsson BO, Hyduke DR. 2013. COBRApy: COnstraints-Based Reconstruction and Analysis for Python. BMC Syst Biol 7:74.

86. Orth JD, Conrad TM, Na J, Lerman JA, Nam H, Feist AM, Palsson BØ. 2011. A comprehensive genome-scale reconstruction of Escherichia coli metabolism--2011. Mol Syst Biol 7:535.

87. Wenk S, Schann K, He H, Rainaldi V, Kim S, Lindner SN, Bar-Even A. 2020. An “energy-auxotroph” Escherichia coli provides an in vivo platform for assessing NADH regeneration systems. Biotechnol Bioeng 117:3422–3434.

88. Seol W, Shatkin AJ. 1992. Escherichia coli alpha-ketoglutarate permease is a constitutively expressed proton symporter. J Biol Chem 267:6409–6413.

89. Nogales J, Mueller J, Gudmundsson S, Canalejo FJ, Duque E, Monk J, Feist AM, Ramos JL, Niu W, Palsson BO. 2020. High-quality genome-scale metabolic modelling of *Pseudomonas putida* highlights its broad metabolic capabilities. Environ Microbiol 22:255–269.

